# Evidence of an Allostatic Response by Intestinal Tissues Following Induction of Joint Inflammation

**DOI:** 10.1101/2024.10.17.618923

**Authors:** Meghan M. Moran, Jun Li, Quan Shen, Sheona P. Drummond, Caroline M. Milner, Anthony J. Day, Ankur Naqib, D. R. Sumner, Anna Plaas

## Abstract

Disrupted intestinal epithelial barrier function has been proposed to be integral to rheumatoid arthritis (RA) progression and pathogenesis. To further define the molecular pathways in synovial inflammation and the response of the intestinal tissues, we have used a rat model of mono-joint inflammatory arthritis, induced by intra-articular injection of Complete Freund’s adjuvant (CFA). The predominant inflammatory response of a single injection of the adjuvant into the knee joint resulted in rapid and reproducible formation of a fibrotic myeloid-infiltrated synovial pannus. Our aim was to determine how intestinal tissues, including the proximal and distal ileum and distal colon, responded to inflammatory changes in the synovium in a temporally coordinated manner by comparing their transcriptomic landscapes using RNASeq analyses. We confirmed the timeline of joint inflammation by knee joint swelling measurement, increased synovial fluid levels of bikunin (a component of both the acute phase protein pre-alpha-inhibitor and inter-alpha-inhibitor) and demonstrated a self-correcting response of trabecular and cortical bone to the CFA challenge. Intestine-specific responses were monitored by 16S microbiome amplicon sequencing, histopathology for mucus layer integrity, and immune cell immunohistochemistry. We present data that shows the intestinal tissue displays an allostatic response to the acute joint inflammation and was region specific. The ileum primarily responded with increased mucus secretion and silencing of T-cell specific pathways, whereas the colon showed a transient upregulation of macrophages, with a broader suppression of immune related and metabolic pathway related transcripts. Interestingly, many neuropathways were activated early but then suppressed later in both the ileum and colon. There were only insignificant changes in the fecal microbiome composition in ileum or colon post-CFA administration. In summary, our data show for the first time a suppression of intestinal inflammatory and immune responses following the induction of joint inflammation and only minimal and transient changes in the microbiome. The results help clarify the molecular responses of intestinal tissues to inflammatory stresses that accompany the pathogenesis of inflammatory joint diseases.

## INTRODUCTION

In the past decade, numerous studies have focused on the role of ‘gut health’ in the progression of several chronic musculoskeletal disorders including rheumatoid arthritis, spondylarthrosis, ankylosing spondylitis, psoriatic arthritis[1, 2], osteoarthritis[3], and bone disorders[4], including osteoporosis[5] and implant loosening[6]. Many of these studies have reported alterations in the composition of the gut microbiota, defined as “dysbiosis”, which includes decreased microbial diversity and/or expansion of specific bacterial taxa that affect the host physiology [7, 8].

However, since acute and chronic inflammatory disease states are accompanied by autocrine and paracrine production of mediators (i.e. cytokines, chemokines, and growth factors), activation of immune cells, and modification of enteric neuronal signals, they can transform the host intestinal environment by altering the structure and the intestinal barrier and function of its resident cells[9, 10]. The intestinal barrier is composed of an outer mucus layer in contact with the commensal gut microbiota[11], anti-microbial proteins, secreted immunoglobulin A molecules[12], a specialized epithelial cell layer[13], and an inner lamina propria that is populated with innate and adaptive immune cells[14]. The multicellular intestinal epithelium consists of multiple cell types, including absorptive enterocytes, mucin-secreting goblet cells, enteroendocrine cells, Paneth cells, and intraepithelial lymphocytes[15]. Furthermore, specific interactions of structural molecules secreted by the epithelial cells, such as mucins and tight junction proteins provide selective permeability for nutrient absorption, while also providing a blockade as primary defense against bacteria to maintain immune homeostasis[16].

Disruption of intestinal epithelial barrier function has been reported as integral to rheumatoid arthritis (RA) progression and pathogenesis [17, 18]. In fact, an early disease response in a collagen induced model for RA (CIA) was shown to be through the Zonulin-CXCR3 mediated dissociation of tight junctions [19]. In addition, elevated HIF2α in intestinal epithelial cells during RA progression can accelerate disease progression. Therapeutic mitigation of those pathogenic responses protected intestinal barrier function and diminished activation of intestinal and lymphatic T-helper 1 (Th1) and Th17 [20]. However, it remains to be determined if the intestinal responses are mediated by a disrupted systemic immune response in RA, and/or if the bone and cartilage destruction mediated by an inflamed synovium (“RA pannus”) is the central link driving the intestinal pathogenesis in RA.

In addition to RA [21, 22], multiple reports suggest the existence of a “joint-gut communication axis” in osteoarthritis, spondylarthrosis, and degenerative disc disease [23–26] that can be part of disease initiation and/or progression. The gut microbiome is proposed as both a target and a mediator in this crosstalk. Likewise an increasing number of studies have explored a correlation between perturbed bone metabolism in osteoporosis, diabetes [27], fracture healing[28], and implant loosening [6] on intestinal homeostasis. These studies also focus predominantly on the role of intestinal dysbiosis and perturbed production of beneficial bacterial metabolites such as short chain fatty acids [21, 29–31], with a few studies implicating the role of T-cells in the communication routes [32, 33].

To further delineate the molecular and cell biological pathways in intestinal responses to joint inflammation, we used a rat model of mono-joint inflammatory arthritis, induced by intra-articular injection (IAI) of Complete Freund’s adjuvant (CFA) [34, 35]. The predominant inflammatory response of a single injection of the adjuvant into the knee joint results in rapid and reproducible formation of a hyperplastic and fibrotic synovial pannus[36] and can produce changes in the intestinal microbiome [37, 38].

The objective of this study was to determine whether inflammatory changes in the CFA-exposed synovium induced temporally related changes in the intestinal barrier of the ileum and/or the colon. The timeline of joint inflammation was assessed by measuring knee joint swelling, joint histology, synovial fluid levels of bikunin (a component of both the acute phase protein pre-alpha-inhibitor (PαI) and inter-α-inhibitor (IαI) [39], bulk-RNA sequencing of synovial tissue, and micro-computed tomography for trabecular and cortical bone changes [40]. At the same time points, intestinal tissue responses were assessed by examining changes in components of the intestinal barrier. This included histochemical analyses of mucus layers, alterations in the intestinal tissue transcriptome using bulk-RNA Sequencing, composition of the mucus layers and abundance and distribution of CD4 and CD8 positive T-cells. 16S RNA analyses for fecal microbiome composition in ileum and colon was also completed.

Taken together, our data show that the intestinal tissue response to an acute joint inflammation was region specific. The ileum responded with increased mucus secretion and redistribution of CD8+ intraepithelial lymphocytes (IEL) as well as a broad suppression of immune function and metabolism-related transcripts. In addition, a concurrent silencing of similar pathways was seen in the colon. The intestinal cell response along with the minimal change in microbiome composition in either intestinal compartment indicated that as a result of an acute inflammation in the joint, intestinal cells will respond with allostasis[41]. This is a mechanism to defend against stress to protect barrier and immune functions as well as microbiota composition and thereby maintain overall function of the gut as a vital organ. This is discussed in the context of examining the mechanisms leading to failure of such preventative responses during progression of chronic inflammatory diseases, such as rheumatoid arthritis, to develop targeted interventions for mitigation of intestinal dysfunction in these diseases.

## RESULTS

### Knee joint responses following IAI of CFA

To establish a time course of knee joint responses following CFA or PBS injections, CFA-IAI and naïve rats were sacrificed at days 3, 7 and 14 post-IAI for analyses (**Fig. S1** and Methods). Knee joint swelling in the medial-lateral orientation had developed by 3d post-CFA-IAI and remained elevated through 14d (p < 0.001, 2-way ANOVA) compared to controls (**Fig. 1A**). There was a significant time effect with CFA-IAI (p = 0.005, 1 way ANOVA) as well as a time x study group interaction (p = 0.008, 2-way ANOVA).

**Figure 1.**
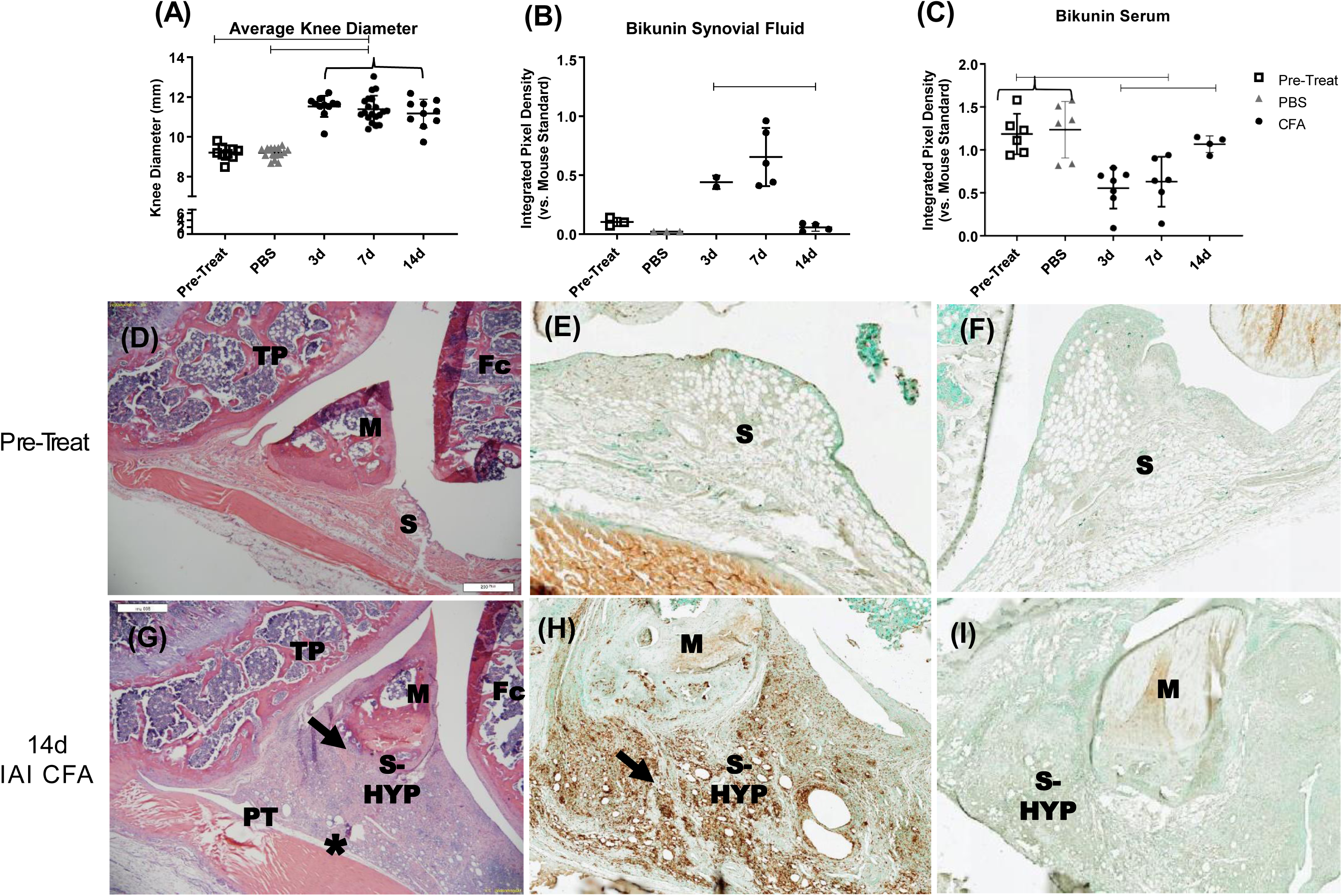
Analyses of knee joint responses to IAI of CFA or PBS. **(A)** At time of sacrifice the diameters of knee joints were measured as described in the Method from rats without treatment (Pre-Treat), after IAI of PBS and 3 days (3d), 7 days (7d) and 14 day (14d) post-CFA. CFA-IAI knees showed significant swelling at all three time points, when compared to pre-treatment or PBS-IAI (Bars= p < 0.001, 1-way ANOVA). **(B,C)** Abundance of bikunin species in synovial fluids (**B**) and sera (**C**) were determined by western blotting as described in the Methods and Supporting Information **Fig. S2**. Note that for pre-treatment (Pre-Treat) and PBS groups, the measurements at 0d and 14d, respectively, were combined since no significant differences were observed between those two groups (general linear model 0d v. 3d v. 14d PBS serum p = 0.226). Coronal FFPE sections of pretreatment and 14d IAI CFA knee joints were stained with H&E (**D** & **G**), anti-CD68 (**E** & **H**) or anti-CD4 (**F** & **I**). The CFA-induced synovial hyperplasia enriched in collagenous ECM and infiltrated by CD68+ macrophages is indicated by black arrows and the swollen patellar tendon by (*). M=meniscus, Fc=femoral condyles, TP=tibial plateau, S=synovium, PT= patellar tendon.

A transient increase in serum effusion into the joint space was confirmed by synovial fluid accumulation of serum-derived bikunin•CS as well as in both inter-α-inhibitor (IαI) and pre-α-inhibitor (PαI) **(Fig. 1B & Fig. S2)** with a concurrent decrease in all these species in the serum (**Fig. 1C**). Furthermore, an abundance of albumin and IgG in synovial fluids at 3d post-CFA-IAI (**Fig. S2**), as determined by SDS-PAGE confirmed the accumulation of serum derived components in the joint space. However, since these proteins were no longer present in the synovial fluids collected at 14d post-CFA, but increased knee diameters persisted, the latter was likely due to joint tissue responses, such as hyperplasia and fibrotic remodeling of the synovium and joint capsule, as well as patellar tendon swelling (**Figs. 1D-I**). No alteration in joint swelling, joint fluid composition or tissue remodeling were noted after PBS-IAI.

### Transient changes in metaphyseal bone following CFA-IAI

As reported by others using the CFA rodent model [34, 42], micro-CT analyses of the distal femoral metaphyseal and epiphyseal bone detected changes in both the cortical and trabecular bone compartments. Surface pitting was observed at 3d post-CFA-IAI in the proximal region of the patellar groove and by 7d the pitting extended to the periosteal surface of the epiphysis and distal metaphysis. (**Fig. 2A**). Adjacent to the periosteal surface, periosteal reactions were present in n=7 7d and n=2 14d rats, primarily in the antero-mediolateral region of the distal femur (representative image, **Figs. 2B & S3**). These reactions occurred transiently coincident with the peak knee swelling, metaphyseal bone loss, and synovial fluid content of bikunin in response to CFA-IAI. By 14d the cortical surface pitting had resolved leading to the smooth appearance of the periosteal and patellar groove surfaces (**Fig. 2B & D**). These cortical changes were associated with significant alterations in cortical bone geometry. Specifically, cortical thickness exhibited significant differences with an overall one-way ANOVA result (p = 0.003), and Bonferroni post-hoc analysis revealed significant changes between 3d and 7d (p = 0.022) as well as between 3d and 14d (p = 0.003). Medullary area also showed an overall significant effect (p = 0.031), with Bonferroni post-hoc significance observed between 3d and 14d (p = 0.041). Similarly, total area demonstrated significant overall changes (p = 0.036), with Bonferroni post-hoc significance between 3d and 14d (p = 0.047). Cortical area showed a trend toward significance (p = 0.069). Trabecular bone volume fraction, normalized to pre-treatment groups, was significantly decreased in the distal femoral metaphysis following CFA-IAI (**Fig. 2E**). This reduction persisted through the 7d peak of inflammation and remained evident at 14d post-CFA-IAI.

**Figure 2.**
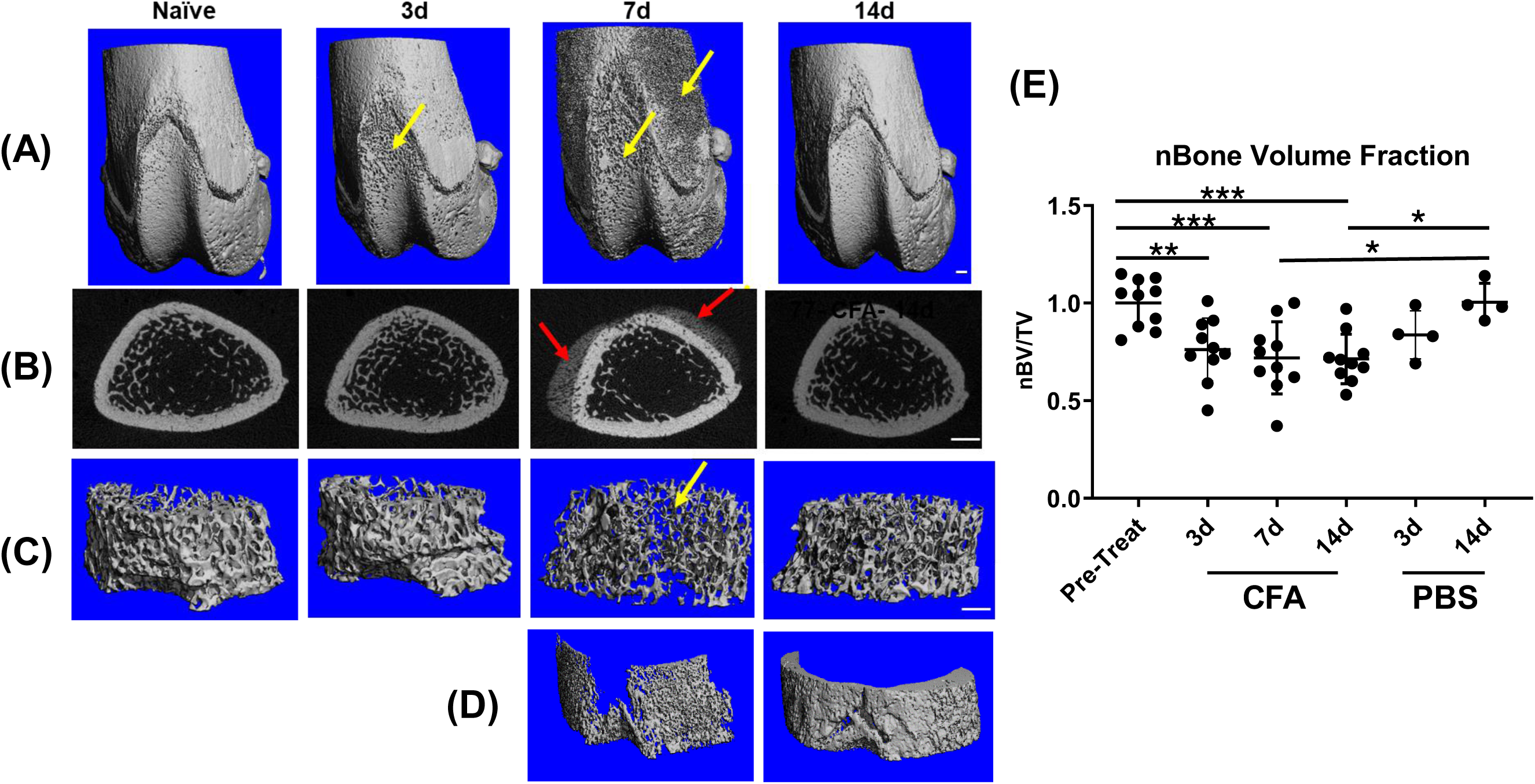
µCT analyses of femoral metaphyses and epiphyses following IAI-CFA. **(A)** 3D rendering of the distal femoral metaphysis and epiphyses show cortical pitting at 3d in the proximal region of the patellar groove with pitting extending to the epiphyseal and metaphyseal regions at 7d (yellow arrows). There is a cortical regenerative response visible by 14d**. (B)** Representative 2D slices through the distal metaphysis adjacent to the patellar groove show the typical appearance of a periosteal mineralization response (red arrows) in the antero-mediolateral region of the femur. This was present after CFA-IAI in (n=7) 7d and (n=2) 14d rats. **(C)** 3D renderings of the metaphyseal trabecular bone adjacent to the cortical bone showed decreased bone volume by 7d post-CFA, which also had recovered by 14d. Scale bars all panels = 1mm. **(D)** Representative 3D renderings of the isolated cortical periosteal mineralization response in the proximal femur at 7d and 14d post-CFA-IAI. The spatial distribution of the cortical pitting at 7d shown in panel A matches the periosteal reaction at 7d and is corrected by 14d post-CFA. **(E)** Bone volume fraction (BV/TV) normalized to pre-treatment group. BV/TV decreases with CFA-IAI compared to both pre-treatment and PBS-IAI. *=p<0.05, **=p< 0.01, ***=p< 0.001.

### Transcriptomic responses of the knee joint synovium following IAI of CFA or PBS

Assessment of changes in tissue- or cell-specific transcriptomic landscapes by RNASeq methodologies has become an essential tool to identify and quantitate responses to disease-causing injurious stimuli in a range of tissues [43]. We have applied bulk-RNASeq methodology and post-sequencing bioinformatic analyses to identify such changes using the Database of Essential Genes (DEG) and associated pathway responses of synovial tissues to IAI of CFA or PBS vehicle control injection.

A global overview of gene expression profiles in synovium at pre- (0d) and post-CFA or PBS is shown in **Fig. 3A**. Whereas PBS injections caused only minimal shifts (increased or decreased abundance) in gene expression profiles at early (3d) and or late (14d) time points, CFA-IAI resulted in extensive changes in the transcriptome, at all time points (3d, 7d, and 14d). DEGs at each post-IAI treatment time point, relative to 0d were calculated and significantly modified genes were outlined in the volcano plots (**Fig. 3B-F**; red dots = activated >1.5 log_2_ FC, p-value<0.05 or suppressed <-1.5 log_2_FC, p<0.05). Using ‘significant fold change’ selection parameters, CFA exposure upregulated transcript abundance for 592, 1037, and 998 genes at 3, 7 and 14d, respectively, but decreased transcripts for 167, 998 and 166 genes at the corresponding time points. By comparison, after sham-IAI-PBS, increases relative to 0d occurred for 32 and 180 genes and suppression for 37 and 67 genes 3d and 14d, respectively.

**Figure 3.**
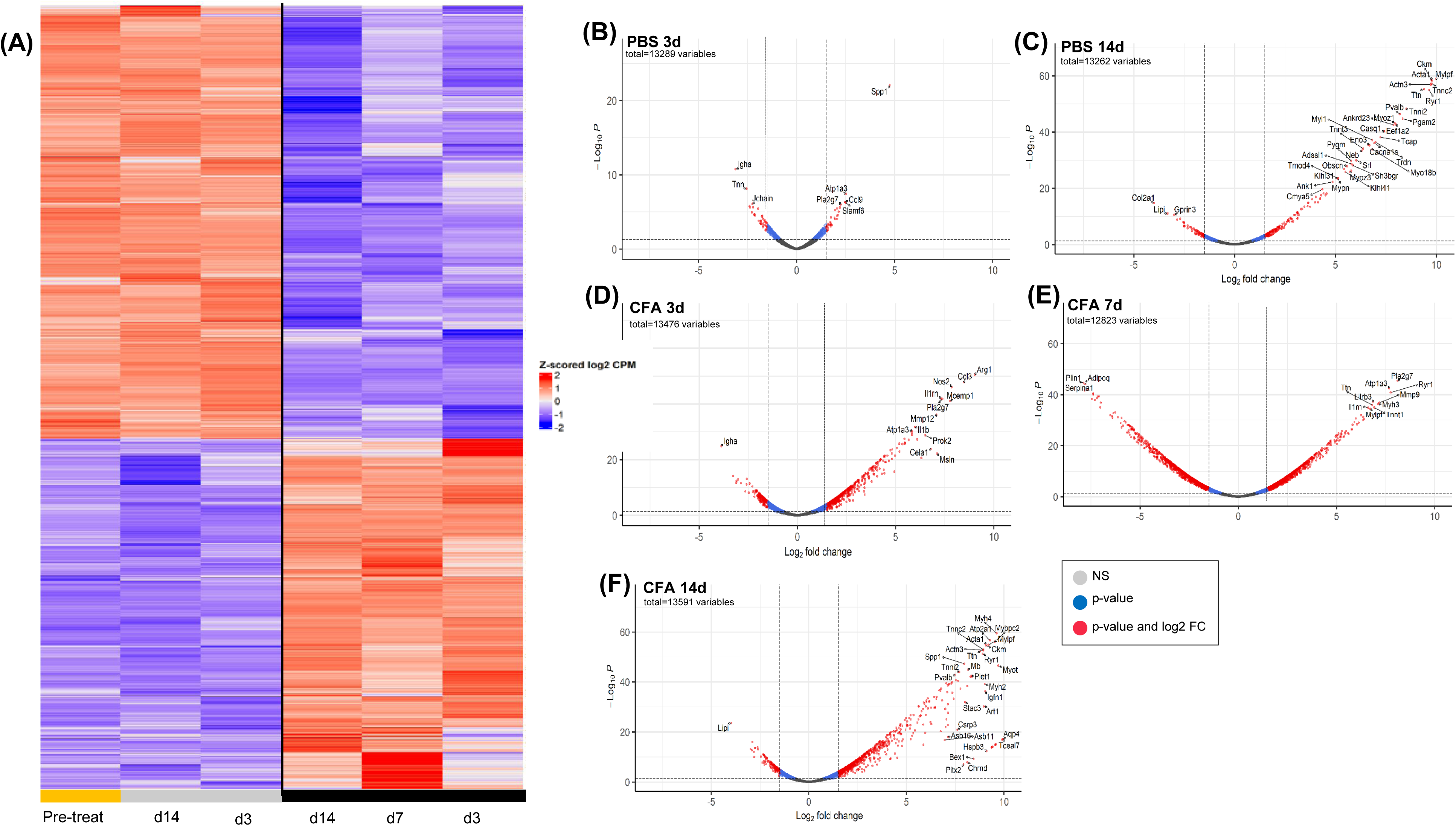
Differentially expressed genes in synovial tissue at 3d, 7d and 14d after IAI of CFA or Saline. **(A)** Heat map illustrating overall shifts in transcriptomic landscape after IAI-PBS (grey bars) or CFA (black bars), relative to d0 (yellow bar). **(B-F)** Statistical significance (p-value) vs. the magnitude of change (log FC) were calculated as described in the Methods and DEGs are summarized in volcano plots. Log FC thresholds of −2 and +2 and an adjusted p-value threshold of 0.05 are delineated by the dotted lines. Upregulated transcripts are on the right, downregulated transcripts are on the left of the lines, and statistically significant DEGs are above the horizontal lines. **Grey dots =** not significant, **Blue dots** = Genes p<0.05, **Red dots=** Genes p < 0.05 and 2-fold change.

The acute (3d) response to CFA-IAI resulted in limited but strong immune and inflammation responses, with >100 fold increases in transcripts of the neutrophil protein *S1009* [44], the macrophage receptor *Clec5a* [45] and the pro-angiogenic protease *Mmp9* [46] as well as the lymphocyte activating factor *Slamf6* [47]. In addition, multiple genes, previously identified with inflammatory arthritis showed sustained modification up to 14d. These included transcripts for pro-inflammatory mediators *Ccl3, Cxcl1, Ccl2, Ccl9,^,^ Nos2, Arg1, Il1m, Il1b,* and *Osm,* their regulators *S100a9* [48], *Slamf6* and *Il1r2*, the C-type lectins *Clec5a* and *Clec4e* [49], as well as *Lcn2* associated with bone damage in RA [50, 51]. In addition, macrophage activating factor *Gdf15* [52], *Adam8* and an integrin, *Itgb8*, from synovial fibroblasts [53] were affected. Notably, transcripts of the *Spp1* gene that encodes the bone protein OPN (osteopontin) and is associated with RA-related synovial inflammation [54], were highly activated in the acute joint response at 3d to both, CFA- and PBS-IAI and remained high up to 7d post-CFA-IAI. Furthermore, transcripts for multiple genes with known functions in innervation and inflammatory pain, *Kcne5, Grin3a, Map2, Erc2, Serpini1, Gprin3, Trhde,* and *Mdga2* were suppressed at all times post-CFA-IAI.

### Transcriptomic responses of the proximal and distal ileum and distal colon following IAI of CFA

Bulk RNASeq analyses was also performed for intestinal tissues collected at 3, 7 and 14d post-CFA-IAI from the proximal and distal ileum and the distal colon. Bioinformatic analyses of sequencing data for DEGs, pathways, and biological process modifications were performed as described in the methods. It should be noted that since PBS-IAI did not yield significant modification of synovial gene transcriptions when compared to CFA exposure (**Fig. 3**), RNASeq analyses of gut tissues was only performed on naïve and CFA-treated gut samples. Gene expression profiles and DEGs for all three intestinal regions (proximal and distal ileum and colon) are shown in **Figs. 4-6**, respectively. These show a wide range of transcriptional changes relative to d0, with statistically significant increases and decreases in DEGs (<log_2_-1.5 and > log_2_+1.5). To supplement the volcano plots in B-D panels, details of DEGs are also shown as heat maps (**Figs. S3A-C** & **S4A-C)**. Together, these data show that changes in affected genes were region-specific and varied with time post-CFA treatment. For example, maximum modifications of the proximal ileal transcriptome were seen at 7d post-CFA, compared to 3d for the colonic transcriptome, with the distal ileum showing a weaker and rather uniform response at all 3 post-CFA time points (**Fig. S3**).

**Figure 4.**
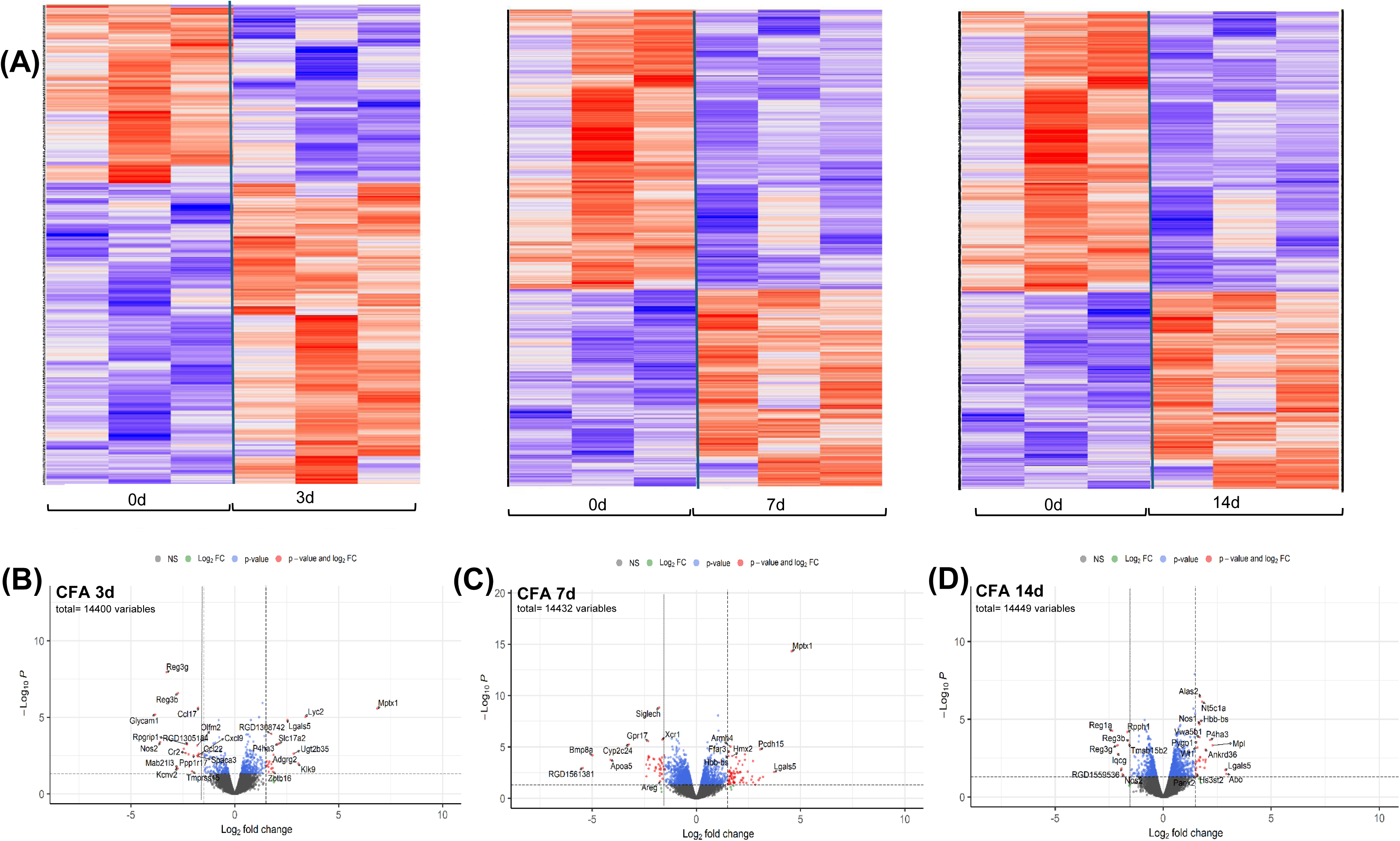
Differentially expressed genes in proximal ileum tissue at 3d, 7d and 14d after IAI of CFA. **(A)** Heat maps illustrating overall shifts in transcriptomic landscape after IAI of CFA relative to d0. Each vertical column corresponds to tissue from a single animal. **(B-D** CFA at 3d, 7 d and 14d respectively) Statistical significance (p value) vs the magnitude of change (log FC) are summarized in volcano plots using the log FC thresholds of −2 and +2 and P-values of 0.05 (dotted vertical and horizontal lines). **Grey dots =** not significant, **Green dots=** Genes 2-fold change, **Blue dots** = Genes p<0.05, **Red dots=** Genes p < 0.05 and 2-fold change.

**Figure 5.**
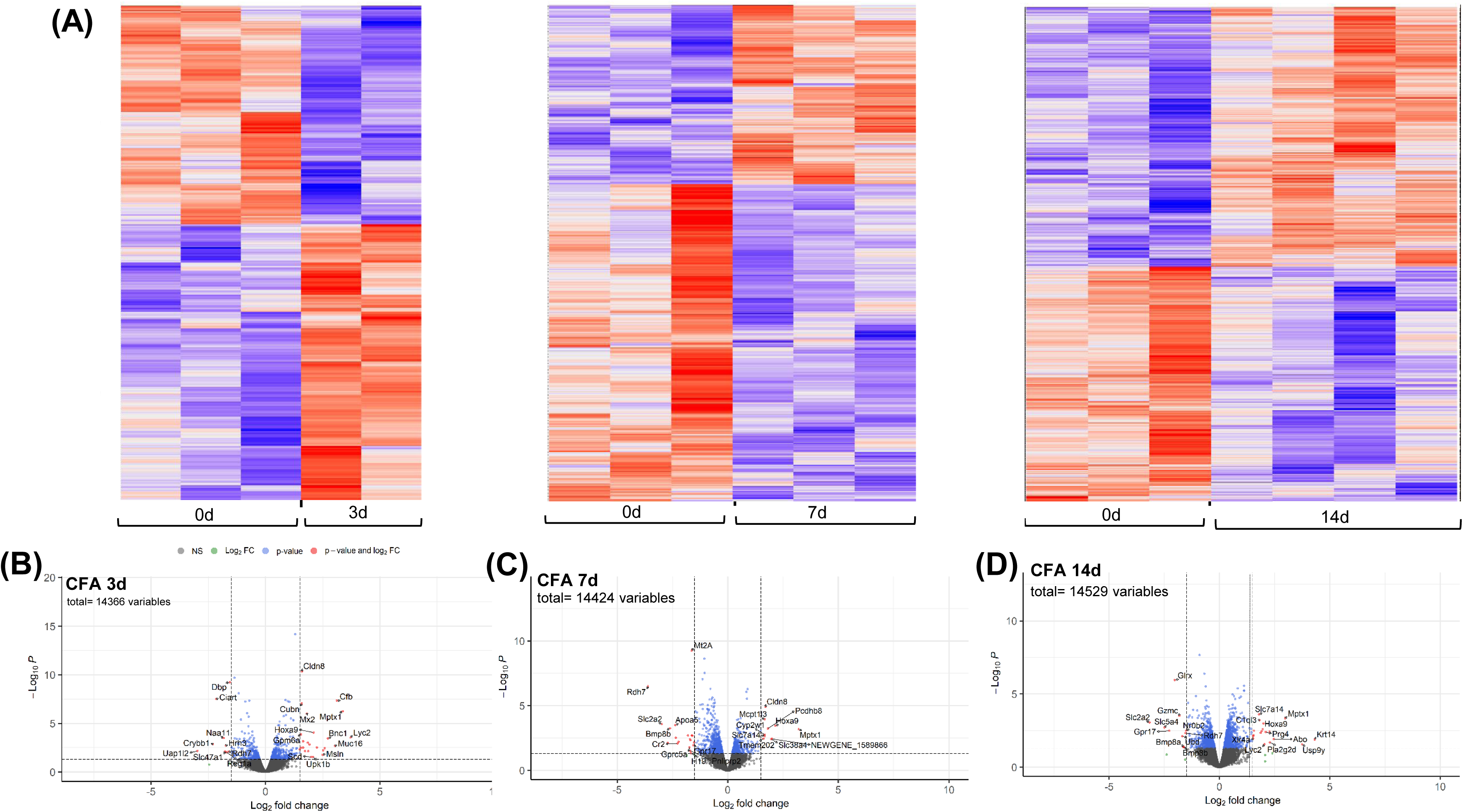
Differentially expressed genes in distal ileum tissue at 3d, 7d and 14d after IAI of CFA. **(A)** Heat maps illustrating overall shifts in transcriptomic landscape after IAI of CFA relative to d0. Each vertical column corresponds to tissue from a single animal. **(B-D** CFA at 3d, 7 d and 14d respectively) Statistical significance (p value) vs the magnitude of change (log FC) are summarized in volcano plots using the log FC thresholds of −2 and +2 and P-values of 0.05 (dotted vertical and horizontal lines). **Grey dots =** not significant, **Green dots=** Genes 2-fold change, **Blue dots** = Genes p<0.05, **Red dots=** Genes p < 0.05 and 2-fold change.

**Figure 6.**
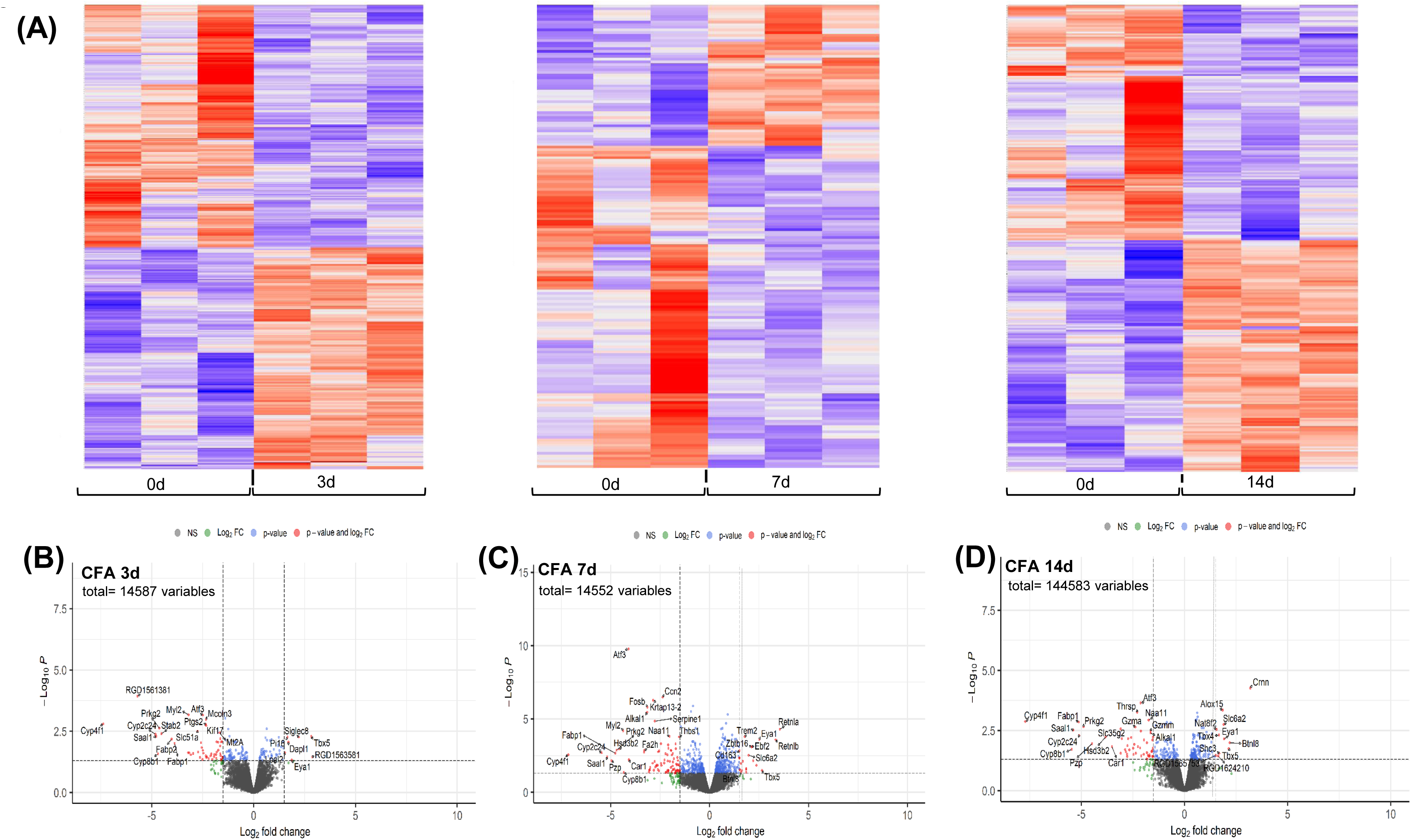
Differentially expressed genes in distal colon tissue at 3d, 7d and 14d after IAI of CFA. **(A)** Heat maps illustrating overall shifts in transcriptomic landscape after IAI of CFA relative to d0. Each vertical column corresponds to tissue from a single animal. **(B-D** CFA at 3d, 7 d and 14d respectively) Statistical significance (p value) vs the magnitude of change (log FC) are summarized in volcano plots using the log FC thresholds of −2 and +2 and P-values of 0.05 (dotted vertical and horizontal lines). **Grey dots =** not significant, **Green dots=** Genes 2-fold change, **Blue dots** = Genes p<0.05, **Red dots=** Genes p < 0.05 and 2-fold change.

Transcripts that were modified at more than one post-CFA time point in each of the three intestinal regions are summarized in **Fig. S4A-C**, and many of these DEGs implicate responses of several cell types in the intestinal barrier. For example, in the proximal ileum, the *Mptx1* pseudo gene [55, 56] is a gut-specific member of the pentraxin family, which is involved in bridging adaptive and innate immunities in inflammatory responses in enterocytes. It is a product of Paneth and epithelial cell transcripts for several REG family proteins, *Reg1a, Reg3b and Reg3G* [57, 58], which are multifunctional molecules secreted by Paneth cells to exert anti-apoptotic, anti-inflammatory and anti-microbial effects were decreased after CFA-IAI. Two genes, *Adgrg2* [59] and sodium and potassium absorption, *Tmprss15* [60], are known to be part of Tuft cell function in the regulation of immune responses. In addition, transcripts for two genes associated with neutrophil responses, *Mpo* [61] *and Mst1* [62], and known to be involved in IBD pathologies, were elevated at 7d and 14d post-CFA-IAI.

In the distal ileum, in addition to the increased transcript for the *Mptx1*, a strong and sustained response of enterocytes was supported by increases in the transcription factor *Hoxa9* [63], the amino acid transporter *Slc7a14* [64], the tight junction protein *Cldn8* [65], the anti-microbial Lysozyme *Lyc2* (*Lyzl-1* [66]), and the *Rdh7* gene, for retinoic acid metabolism [67]. Downregulated expression was seen for epithelial cell specific genes, involved in glucose transport and uptake, *Gpr17* [68], *Slc2a2* [69] and *Slc5a4* (*Sglt3*) [70].

The most profound transcriptional response to CFA-IAI was seen in the distal colon with decreased mRNA levels for a wide range of genes (**Figs. S3C** & **S4C**). Downregulated transcripts were seen for genes involved in regulation of intestinal immune responses, including the immune cell infiltration-related *Serpine1* [71] and the transcription factor *Atf3* [72, 73]. Several of the modified transcripts also point to cell-specific responses in the colon including the epithelial cell-associated genes *Btnl8* [74], *Fabp1* [75], *Prkg2* [76]*, Car1* [77] and *Dmbt1*[78]; the fibroblast/myofibroblast associated genes *Cxcl11* [79], *Nr4a1* [80], and *Tbx5* [81]; as well as the enteroendocrine cell gene *GcG* [82]. Small, but significant increases were also seen for the transcription factor *EyA1* [83], which is implicated in the activation of cell proliferation and EMT in colon cancers.

### Pathway and biological process allocation of DEGs using GO enrichment

To assign a functional significance of modified gene transcripts within the context of cellular pathways in synovium and intestinal tissues, GO enrichment analyses was performed using the KEGG pathway database. Individual pathways were further assigned to the following six biological processes listed in that database: Metabolism, Cellular Processes, Genetic Information Processing, Environmental Processing, Immune System and Nervous System. Note that all gene sets with a log FC value of <-0.3 and >+0.3, and an adjusted p-value <0.05, were included. The data are displayed in dot plots for synovium (**Figs. S5** & **S6**), proximal ileum (**Fig. S7**), distal ileum (**Fig. S8**) and distal colon (**Fig. S9**) and are also summarized in Venn diagrams (**Figs. 7** & **8**) Following CFA-IAI, a large number of pathways were activated in synovial tissue (**Figs. 7A** & **S5A**) (i.e., 45, 40 and 37 at 3d, 7d and 14d, respectively) or suppressed (i.e. 22, 32 and 6 at 3d, 7d and 14d, respectively). As expected from previous reports of the CFA model [34, 36], the adjuvant produced robust and persistent immune and inflammatory reactions in the synovium. Activated pathways included “NK cell mediated cytotoxicity”, “Neutrophil extracellular trap formation”, “Th1 and Th2 cell differentiation”, “Th17 cell differentiation” and “B cell receptor signaling” pathways at 3d, 7d, and 14d post-CFA-IAI. This was accompanied by a stimulation of “Toll-like-receptor”-, “C-type lectin-” “IL17-” and “NOD-like”-immune cell signaling characteristic of immune cells. Other inflammation-related pathways such as of “cytokine–cytokine receptor interactions”, “chemokine signaling” and “NF-kB signal transduction” were also stimulated. Throughout the post-CFA period examined, a range of other pathways were suppressed, such as the Hippo pathway [84, 85], which can lead to induction of cell division and the suppression of apoptosis by PPAR signaling. This would be expected to interfere with control of the inflammatory response to TNF-α in synovial cells [86]. Other notable adjuvant-activated responses were seen in lysosomal, phagosomal, apoptotic, and necroptotic pathways, while neuronal system pathways such as “axon guidance” and “glutamatergic synapse function” were suppressed. By comparison, PBS-IAI resulted in a very minimal response in the synovium. The only modified pathways post-PBS-IAI that showed overlap with those modified by CFA-IAI were “heparan sulfate biosynthesis”, “RNA polymerase”, and “tryptophan metabolism” at 3 d and “neuroactive ligand-receptor interaction” and “HIF-1 and calcium signaling” pathways at 14d. There was a lack of immune or inflammation related pathways changed by PBS. This is in contrast to what was expected and previously shown as a mild sham-IAI response in the clinical setting for sham injection[87] or joint space lavage[88].

**Figure 7.**
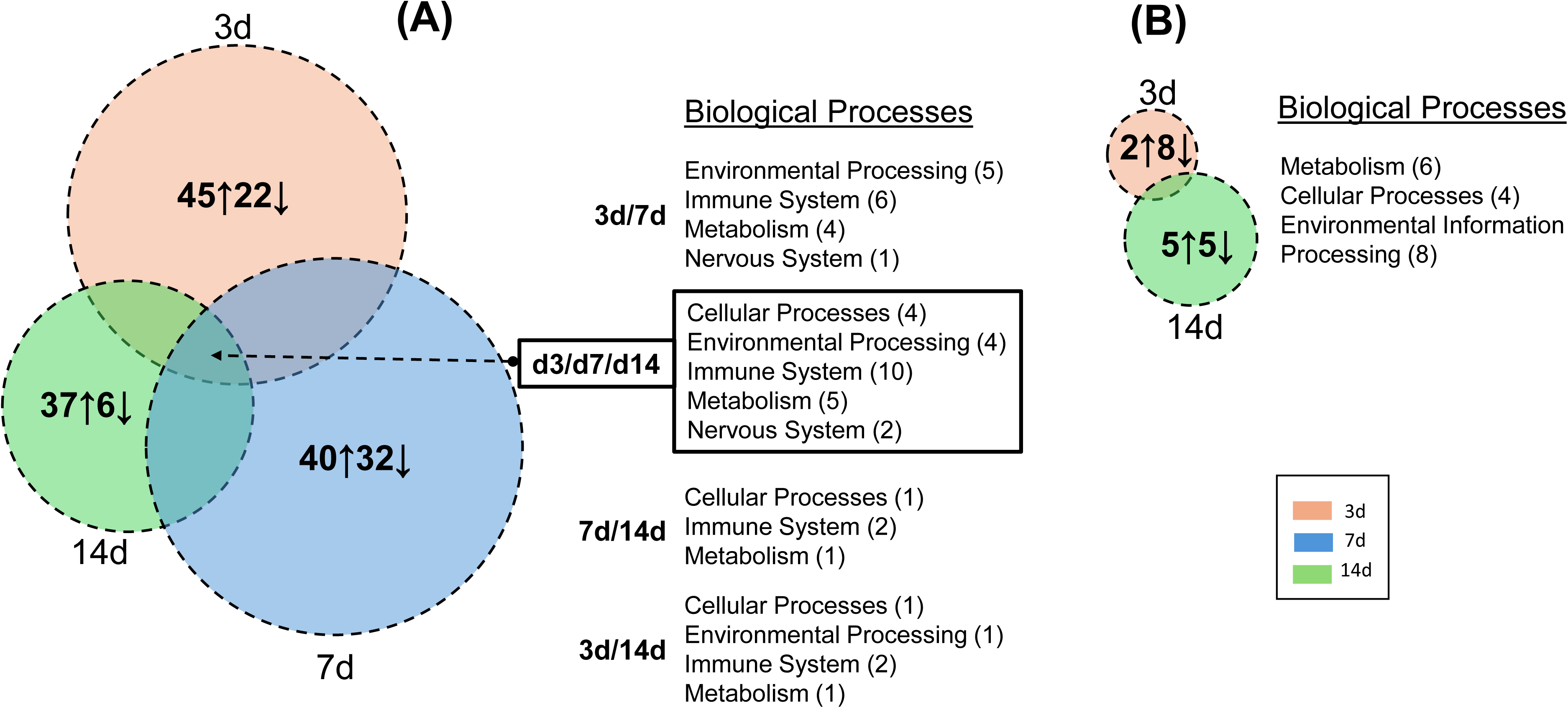
Venn diagram illustration of modified biological process in synovium. Following **(A)** CFA or **(B)** PBS exposure. The numbers in individual circles indicate the total pathways activated (↑) or suppressed (↓) at each time point, and these are individually displayed and in color-coded KEGG-based grouping of “Biological Processes”. Processes modified at two (3d/7d, 7d/14d, 3d/14d) or three (3d/7d/14d) times points are listed to the right of the Venn diagram and the number of affected pathways in each Process is shown in brackets.

**Figure 8.**
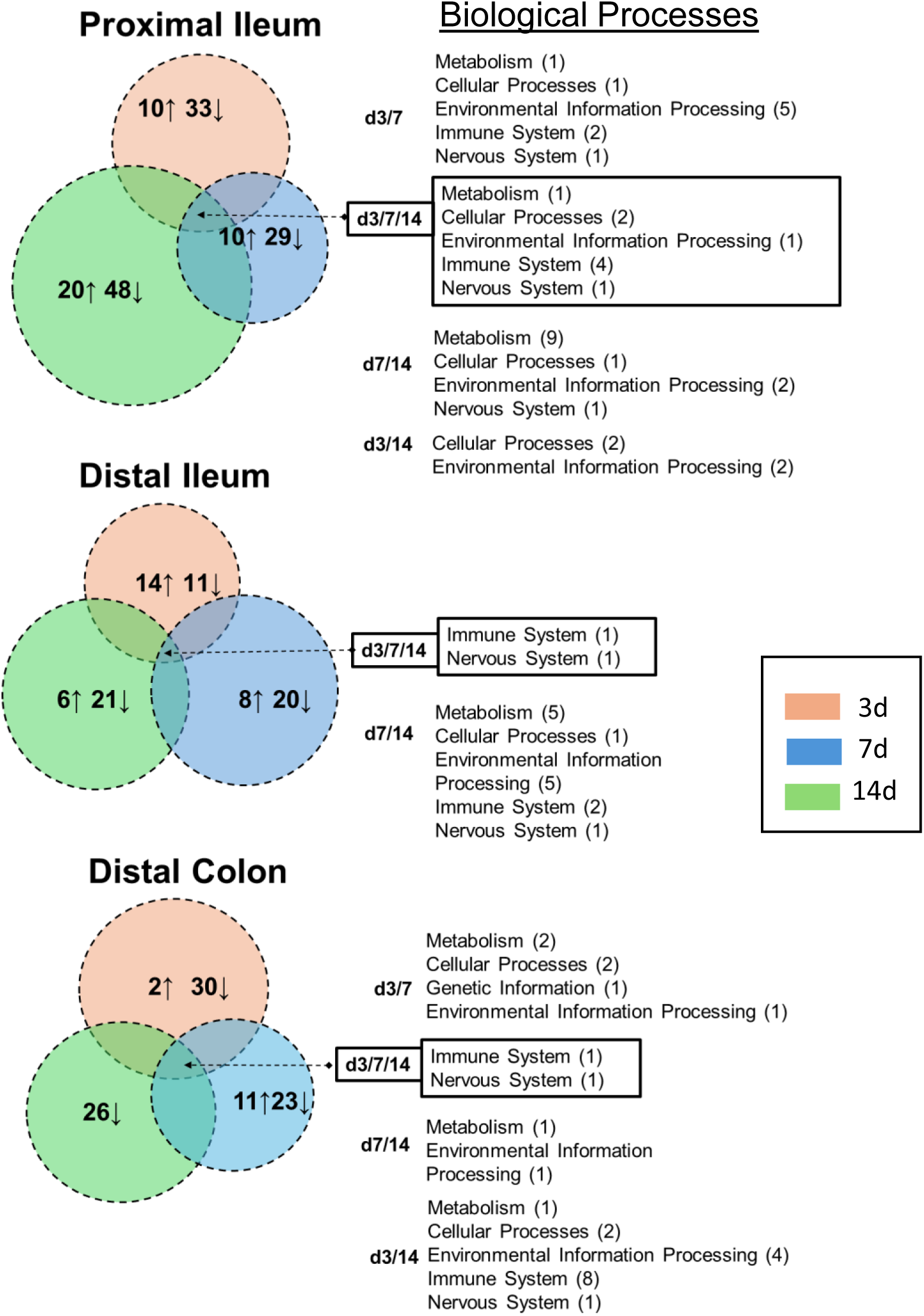
Venn diagram illustration of modified biological processes in proximal and distal ileum and colon following CFA-IAI. Numbers in individual circles indicate the total pathways activated (↑) or suppressed (↓) at each time point, and these are individually displayed and in color-coded KEGG-based grouping of “Biological Processes”. Processes modified at two (3d/7d, 7d/14d, 3d/14d) or three (3d/7d/14d) times points are listed to the right of the Venn diagram and the number of affected pathways in each Process is shown in brackets.

The corresponding GO analyses of intestinal tissues **(Figs. 8 & S6-S8)** showed fewer affected pathways than those identified in the synovium. As expected from the extensively downregulated transcriptome (**Figs. 4-6**) after CFA-IAI, many of the intestinal pathways were suppressed, most notably those associated with the immune system. In the proximal ileum, “Th1 and Th2 cell differentiation”, “Natural killer cell mediated cytotoxicity”, “Th17 cell differentiation”, and “Antigen processing and presentation” were suppressed over the entire experimental timeline. In the distal ileum, the “Intestinal immune network for IgA production” and the “Antigen processing and presentation” pathways were suppressed at 7d and 14d. These immune pathways were also suppressed in the distal colon, together with “B cell receptor” and “T cell receptor-signaling, and “Leukocyte trans endothelial migration”. Transcripts for a range of metabolic pathways and signaling pathways were also decreased in both the ileum and colon.

A comparison of pathway-specific responses between synovial and intestinal tissues, is shown in **Fig. 9**. Although many of the same pathways were affected, activation responses in the synovium gave corresponding suppression responses in the intestinal tissues, and this was particularly evident during the peak joint inflammation responses at 3d and 7d. This, together with the DEG data, confirms a generalized ‘silencing’ of inflammatory and immune reactions in the intestine post-CFA-IAI. This likely constitutes a shielding response to maintain intestinal barrier function in the presence of an acute innate immune and inflammatory insult at a remote tissue site.

**Figure 9.**
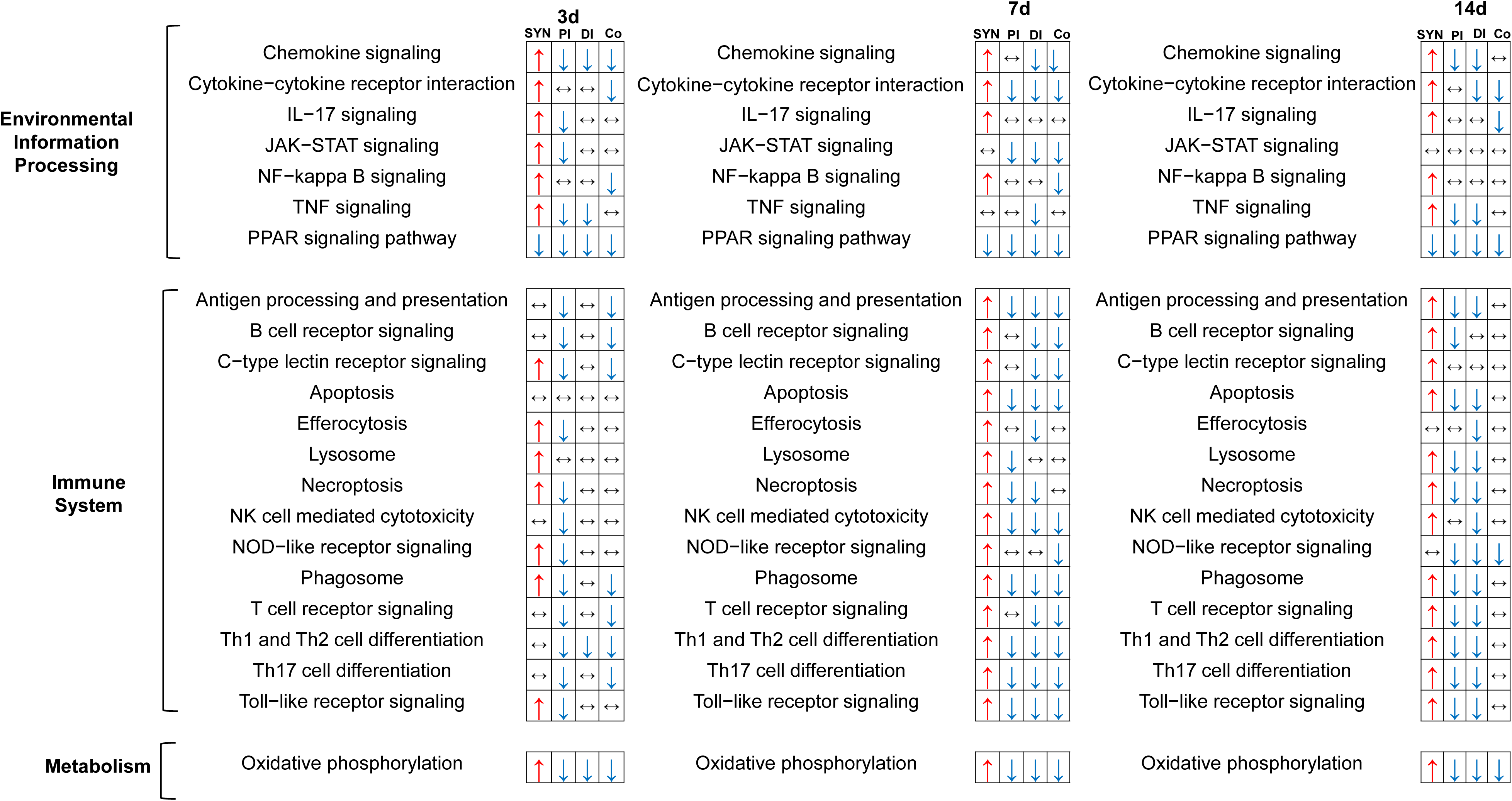
Comparisons of modulated pathways in synovial and intestinal tissue following CFA-IAI. Activated pathways are marked with a red upward pointing arrow (↑) and suppressed pathways with a blue downward pointing arrow (↓). No change is indicated by a horizontal double arrow (↔). SYN= synovium, PI = proximal ileum, DI = distal ileum, Co= Colon.

### Effect of IAI CFA on intestinal immune cells and mucin metabolism

To test for alterations in intestinal immune cell population, such as macrophages and T-cells, we performed IHC analyses of sections from Swiss-rolled ileum and colon using anti-CD68 for macrophages, anti-CD4 and anti-CD8 for resident T cells (**Figs. 10A-C** and **S10-S12**). In healthy tissue, CD68+ macrophages were almost exclusively located in the lamina propria of the ileum and the colon, and there was no change in their abundance or localization along the ileum. However, in the colon, at 3d post-CFA-IAI, transient increases in CD68+ve cells surrounding crypts and interspersed between enterocytes along the villi was seen (black arrows, **Fig. 10A**) was seen. This acute buildup of macrophages in the colon was further supported by increased transcript abundance of macrophage-specific genes *Trem2*[89] and *C1qc*[90] (**Fig. 10D**).

**Figure 10.**
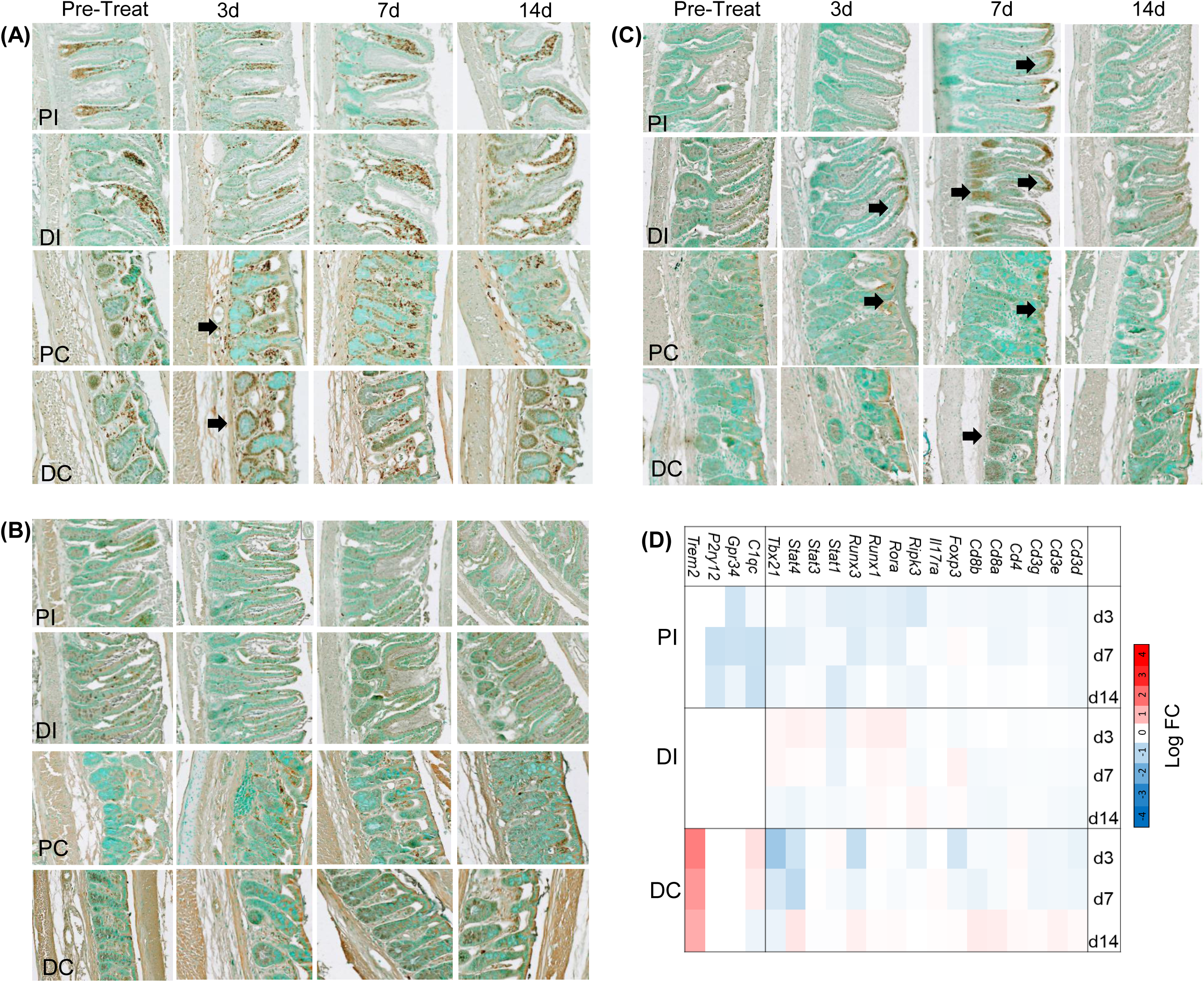
Effect of CFA-IAI on macrophage and T cell responses in distal Ileum and colon. Adjacent FFPE-sections from proximal and distal regions of the ileum and colon were stained with (**A**) anti-CD68 to localize macrophages or (**B**) anti-CD4 and (**C**) anti-CD8 to localize T-cells. Black arrows indicate regions of increased reactivity post CFA. (**D**) DEGs typically associated with intestinal macrophages or T cells caused by CFA-IAI are summarized in the heat map. PI=Proximal Ileum, DI= Distal Ileum, PC= Proximal Colon, DC= Distal Colon.

CD4+ T cells were distributed along the villi in the ileum and additionally in the apical regions in the colon, likely expressed by the intraepithelial lymphocytes[15] (IELs), which were unchanged during the post-CFA period (**Fig. 10B**). In naïve intestinal tissue, the anti-CD8 also stained predominantly IELs **(Fig. 10C)**. However, at 3d and 7d post-CFA-IAI, increased staining of such cells in the apical part of the villi was seen in many regions throughout the ileum and the colon. In addition, at 7d CD8+ve populations had also accumulated in the crypts (**black arrows**, **Fig. 10C**). By 14d, however, the abundance of these T-cells and their locations had returned to those seen in naïve intestinal tissues.

In keeping with the protective response of intestinal tissues to knee joint inflammation, there were no changes in cell proliferation (Ki67 staining) in any intestinal region (**Fig. S13**). We also observed increased secretion of mucins in the distal ileum, during the acute response phase at 3d & 7d, shown by Alcian blue staining (**Fig. 11A**), and supported by increases in transcript abundance for the gel-forming mucins *Muc2* and *Muc5b*[91] (**Fig. 11C**). Furthermore, histochemical staining with O-linked oligosaccharide specific lectins (GSL I, MAL II, and UEA) showed an increased reactivity in mucus with the GSL I (specific for terminal α-N-acetylgalactosamine or α-N-galactose residues)[92] in the distal ileum at 3d post-CFA-IAI (**Fig. 10B**). There were no detectable changes in staining with MAL II (terminal α-2,3 sialic acid) or UEA (α-linked fucose) (**Fig. S14**). Modulation of mucin glycosylation in response to the joint inflammation was further underlined by altered mRNA transcripts for multiple glycosyltransferases, including *St6galnac1* (**Fig. 10C**), the major sialyltransferase in goblet cells, known to be induced by microbial pathogens and critically important for maintaining mucus integrity [93].

**Figure 11.**
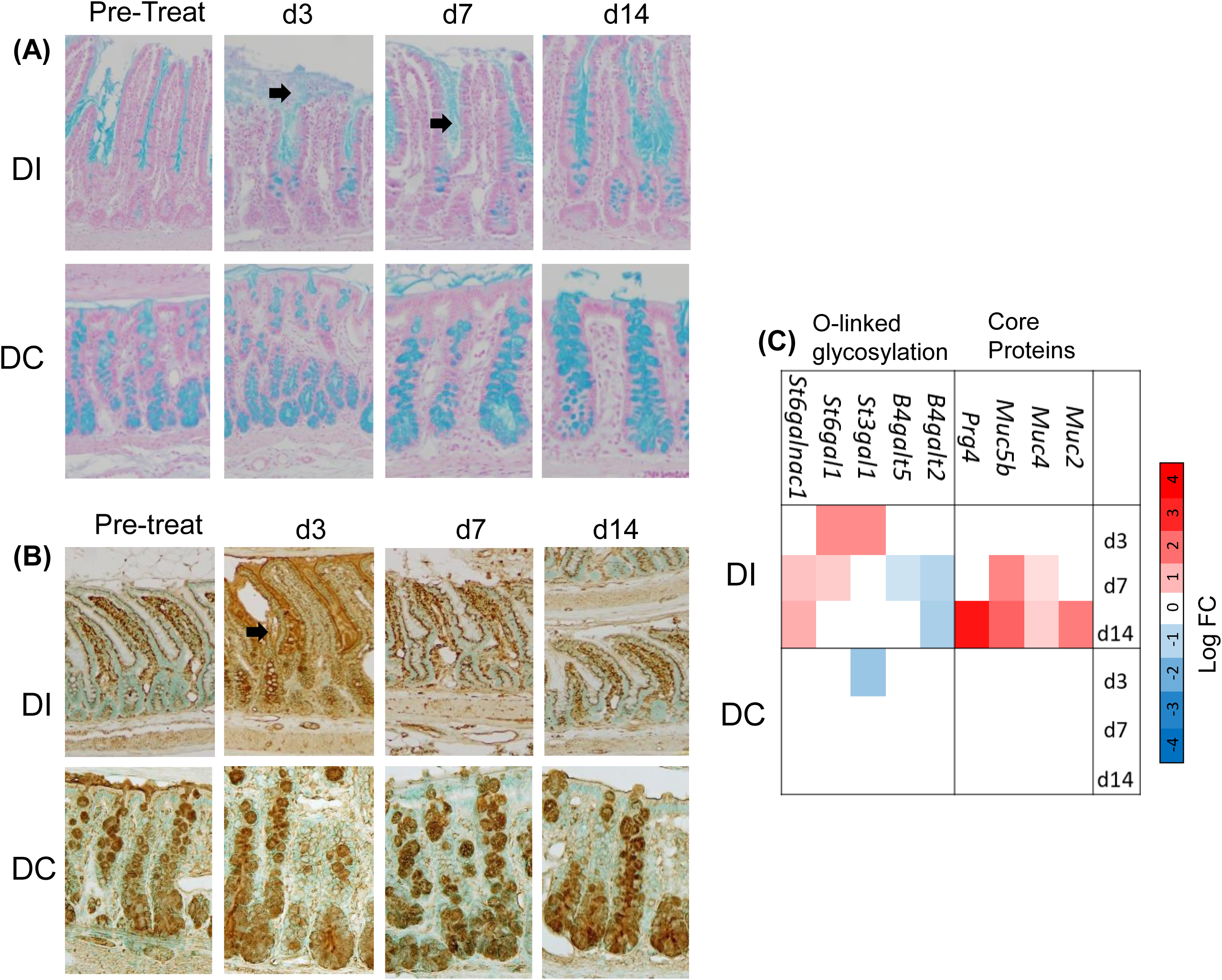
Modified mucin secretion in distal ileum and colon following CFA-IAI. Adjacent FFPE-sections of Swiss-rolled ileum or colon stained with Alcian Blue (**A**) or the biotinylated GSI lectin (**B**) as described in the Methods. Brown= positive stain; Black arrows show the increased mucin secretion with enhanced GSI staining in the DI at 3d post IAI of CFA. Modified gene transcripts in the mucin biosynthesis pathways in proximal and distal regions of the ileum and the distal colon, which were affected by IAI of CFA are summarized in the heat map in panel (**C).** PI=Proximal Ileum, DI= Distal Ileum, PC= Proximal Colon, DC= Distal Colon.

### Effect of CFA-IAI on Digestive Functions and Microbiome Composition

Based on our RNASeq data, several digestive system pathways throughout the different regions of the intestine were affected by the inflammatory response of the knee joint. For example, transcripts encoded by genes associated with ‘mineral absorption’ were modulated in all 3 regions of intestine and were largely decreased compared to pre-treatment. The distal colon showed suppression of the fat digestion pathways, with decreases in expression of 4 associated genes **(Fig. 11A).**

Notably, there was no evidence for development of dysbiosis in the model. The fecal microbiota from the distal ileum and colon were minimally altered in response to the knee joint inflammation and respective intestinal cell responses. Alpha diversity for within-group comparison was unaltered, as shown by evenness (p = 0.697), Shannon index (p = 0.997), and Simpson index (p = 0.998). Relative bacterial abundance (>1%) for the distal ileum and the colon (**Fig. 11B**) at the phylum level showed a slight shift from Naïve to 3d, 7d and 14d post-CFA-IAI. Amongst this shift, there was a decrease in *Romboustia* in the ileum at 3d post-CFA, followed by an increase at 7d and a return to naïve levels by 14d. *Lactobacilus* showed the opposite shift with an increase by 3d, decrease by 7d and a return to naïve levels by 14d. This change in abundance supports a mild and importantly, a limited transient response of the microbiome when acute inflammation in the joint peaks at 7d post-CFA-IAI. Due to the observation of a transient change in mucus layer 16S data were mined for the abundance of mucin-degrading bacteria (*Akkermansia, Alistipes*) and sialic acid cleaving bacteria (*Bifidobacterium*) (**Figs. 12** & **S14**). Apart from the appearance of a low levels of *Akkermansia* in the feces from distal ileum and distal colon post IAI of CFA, the joint inflammation had not detectable effect on the other two species.

**Figure 12.**
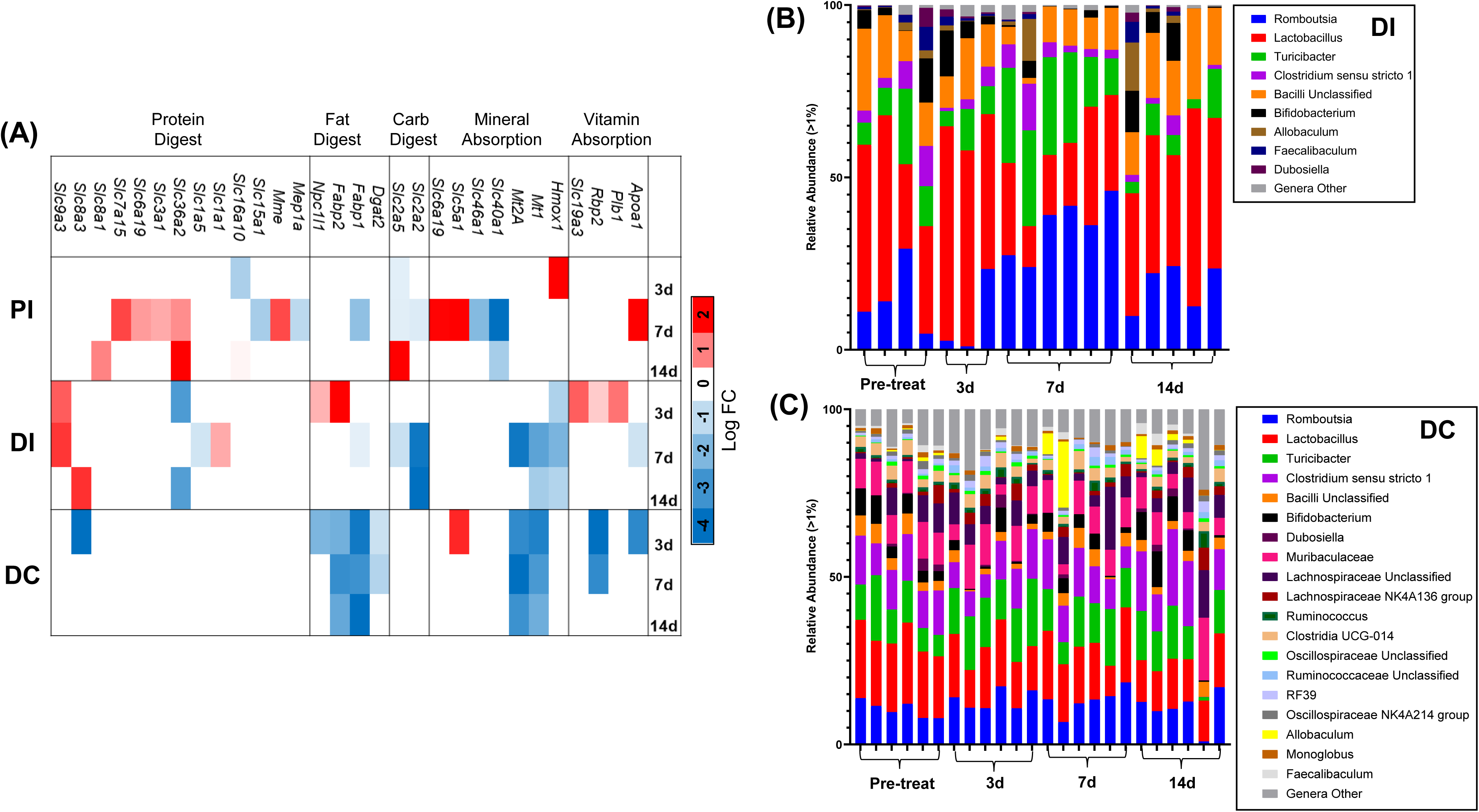
Effects of CFA-IAI on genes in associated with digestive functions and microbiome compositions in distal ileum and colon. **(A)** Data were obtained from the DEGs from analyses of RNASeq data (see Figs. 3 & 5) and are shown as log fold changes (p<0.05) as a heat map. **(B** & **C)** 16S Amplicon Sequencing of Fecal Microbiome was performed as described in the Methods. Relative bacterial abundance (>1%) is shown for the distal ileum (DI, **B**) and the distal colon (DC, **C**) at pre-treatment, 3d, 7d and 14d at the phylum level. Individual phyla are color-coded and listed on the right.

## DISCUSSION

In this study we present novel data on coordinated cell and molecular biological changes in the synovium and intestinal tissues in the acute CFA-induced knee joint inflammation model. Due to its reproducibility and rapid inflammation responses in the injected joint, this model has been used extensively for preclinical efficacy testing of a wide range of anti-inflammatory [94, 95] and analgesic [96–99] RA drugs. A number of these studies have chosen the inflamed synovium as a target, with a focus on the cytokines and the NFĸB signaling pathway [100–103]. This model has also been used in several studies to examine the effect of joint inflammation on the gut microbiota [37, 96] supporting the existence of a “cross-talk” between joint and remote tissues like the gut. However, only limited data of the cell biological responses in the intestinal barrier tissues and innate immune cell populations are currently available. Our data reported here provide a detailed and comprehensive assessment of cellular and molecular biological changes underlying inflammatory responses during the formation of a synovial pannus and corresponding intestinal responses.

Our data confirm an early and robust response of the joint to the adjuvant at 3d and 7d post-CFA-IAI. In addition to measurements of joint swelling, we showed joint effusion by elevated levels of serum albumin and serum-derived bikunin-containing complexes[39] in the synovial fluids. The latter include the liver-derived IαI and the acute phase proteoglycan PαI, which are known to accumulate in synovial fluids in osteoarthritis and rheumatoid arthritis[104–106]. In this context, it is notable that in an inflammatory tissue environment, bikunin-containing complexes can act as donors of their chondroitin sulfate bound-heavy chains for transfer to hyaluronan in the presence of TSG6[107–110]. This results in a crosslinked extracellular hyaluronan network[111] that is permissive for leukocyte cell adhesion[112, 113]. Generation of such a matrix in joint tissues is likely involved in the acute phase of this model, since RNASeq data show a ∼2-fold increase in mRNA levels for TSG6 (*Tnfaip6*) as well as hyaluronan synthase 2 (*Has2*), at 3d post CFA, with both genes returning to pre-inflammation levels by 7d (data not shown). Moreover, following the acute inflammatory response, the perimeniscal synovium was transformed into a hyperplastic tissue with a fibrotic matrix that was extensively populated by myeloid cells, but no significant presence of T-lymphocytes was detected.

Recent studies of RA patient-derived synovial biopsies have identified subgroups of synovial “pathotypes” [114, 115]. Interestingly, our data suggest that CFA-IAI results in formation of the mixed myeloid/fibroid, but not the lymphoid pathotype. The former is reported to be treatment-resistant to many current DMARDs contributing to persistent “low joint disease activity” of many RA patients. This underscores the utility of this model to develop novel therapeutic approaches targeting joint tissue destruction in inflammatory arthritis. The development of a macrophage-rich synovial tissue is supported by the RNASeq based pathway analyses. Firstly, post-CFA activation of signaling pathways, including Notch, JAK/STAT, NF-κB, and MAPK [116], metabolic reprogramming, and increased glycolytic pathways are known to control macrophage polarization towards the pro-inflammatory phenotype in RA. In addition, biosynthetic pathways for arginine, a substrate for the inducible nitric oxide synthase (iNOS) in macrophages, indicates elevated production of nitric oxide (NO) and citrulline, both of which are key metabolites in the inflammatory pathogenesis of RA [117, 118]. Notably, persistent activation of the “lipolysis in adipocytes", fatty acid metabolism, as well as cortisol synthesis and secretion pathways seen in our studies have also been reported to be a markers for a highly pro-inflammatory synovial subtype in OA and RA patients[119–121]; and is consistent with turnover of the sub-synovial adipose tissues induced by joint inflammation.

The development of a pro-inflammatory macrophage-rich synovium is also likely responsible for epiphyseal and metaphyseal periosteal surface pitting present at 3d and 7d post-CFA, in regions that were invaded by the hyper-proliferative pannus[122]. A transient downregulation was present of propanoate, a short-chain fatty acid known for its role in anti-inflammatory macrophage polarization, and specifically in suppression of osteoclastic activities in particle-induced aseptic osteolysis[123–125]. Re-establishment of a smooth periosteal surface by 14d supports the conclusion that the specific macrophage-mediated inflammatory joint tissue responses in the CFA-model are self-limiting, which is further supported by a robust increase in transcripts for oxidative phosphorylation[126]. This pathway plays a critical role during the polarization of the macrophage phenotype from a pro-inflammatory, glycolysis-dependent type to the anti-inflammatory type[127, 128]. This together with a mineralization response seen in the post-CFA reactive periosteum would account for the reversal of periosteal surface damage. On the other hand, trabecular bone recovery did not occur in keeping with published data[129], as remodeling cycles in this compartment are long compared to the 14d time frame of our experiment with timing of osteoclast recruitment to this site, lifespan of osteoclasts, and their lifespan in rats is on the order of months[130].

On the other hand, there was no attenuation of the increases in cytokine and chemokine signaling pathways or activated T and B cell characteristic pathways, suggesting that even though several inflammatory responses in the joint are transient, many of the CFA-induced cellular and metabolic changes are in keeping with an irreversible modification of the synovial tissues. Moreover, additional challenges to such joints could lead to more severe innate responses and eventually lead to activation of the adaptive immune system and development of a chronic inflammatory arthritis. It should also be noted that only a few DEGs during intestinal responses to CFA-IAI overlapped with those detected for the synovium. We show that *Nos2, S100a9, Cxcl1, Ccl2, and Lcn2* increased in the synovium and decreased in the intestine, whereas *C1ql3* and *Serpine1a* were decreased in the synovium and increased in the intestine.

Gross evaluation by histological assessment of the ileum and colon revealed no detectable changes in the overall appearance of the tissue during the 2-week post-CFA period. This included intactness of intestinal villi, no increases in vascularization of the submucosa (data not shown), and unchanged cell proliferation indices, as determined by distribution and abundance of Ki67+ cells. There were no marked changes in the abundance of goblet cells throughout the ileum and colon, with only a transient change in mucus secretion in the distal ileum. In addition, the microbiota composition was minimally affected, with only a mild and transient response during peak joint inflammation (7d post-CFA-IAI) and normalized by the 14d time point. Moreover, using immunohistochemistry, we detected only transient alterations in immune cell abundance, including increased CD68+ macrophages in crypts of the colon and increased CD8+ T cells at the apical surface of villi in the ileum. Moreover, such changes occurred in sporadic regions of the intestinal compartments.

Our data contrast with the widespread intestinal manifestations observed in the chronic rodent models of inflammatory arthritis induced by immunization with type 2 collagen[131]. In fact, our bulk RNASeq data analyses from the proximal and distal ileum and the distal colon indicate a widespread "allostatic" response[41] to the CFA-induced inflammation and pain stressors in the joint. We observed significant suppression of transcriptomes compared to levels seen in those tissues pretreatment and included downregulation of genes in multiple pathways controlling pro-inflammatory signaling, immune activation, and cellular metabolism. Such broad “gene silencing” might be expected as a response to prevent chronic damage to the essential functions of the intestine, including nutrient digestion and absorption, as well as defense against bacterial invasion and support microbial symbiosis. In particular, our data showed containment of cytokine and chemokine signaling and inhibition of the oxidative phosphorylation (OXPHOS) and PPAR-ү pathways throughout the post-CFA-IAI period. OXPHOS and mitochondrial function are fundamental for metabolic homeostasis of all cell types of the intestine and imperative to a functional barrier, however, enhanced flux through this pathway occurs when increased ATP is needed by immune cell during their activation and cytokine production. Furthermore, mitigation of reactive oxygen species (ROS), generated as by-products of enhanced OXPHOS, would prevent their damaging effect on intestinal function [132], including disrupted barrier function, dysbiosis, and impaired immune responses [133, 134].

Genes in a second pleiotropic signaling pathway, the PPAR-ү pathway, extensively studied in colon cancers[135] were also downregulated in the post-CFA-IAI period, and this occurred both in the intestinal tissues and the synovium. In the context of the latter, this activity is pivotal in regulation of adipose tissue metabolisms, and its downregulation is a hallmark of inflammatory responses [136], which is consistent with loss of adipocytes in the myeloid/fibrotic induced synovium from CFA-IAI. A protective effect of its downregulation in the intestinal tissues might be attributed to silencing its cross-signaling with a range of other signaling pathways, including NFκB, Notch [137], JAK/STAT [138, 139], MAPK [140], and Wtn/β-catenin [136]; some of which were also found to be downregulated in the post-CFA intestines. In summary, we provide, for the first time, data on temporally coordinated cellular responses between joint tissues and the intestine following an acute inflammatory insult by ICFA-IAI into the knee joints of skeletally mature rats. Additional application of spatial transcriptomics to depict anatomical location of transcriptomic alterations in the intestinal barrier, together with ATACSeq[141] to identify regions of the genome that are accessible or blocked would further clarify regulatory mechanisms that could serve as therapeutic targets locally or systemically, during inter-organ communication in chronic arthritis.

Transmission of signals between diseased tissues and the gut has been postulated for a wide range of disease conditions, but the molecular and cell biological mechanisms remain largely undefined, except for the much-studied bidirectional pathway between the brain and the intestine[142], which includes cytokine, chemokines, migrating or circulating immune cells, and microbiota-derived metabolites. More relevant as a potential mediator of intestinal responses to joint inflammation is the enteric nervous system (ENS), which is a highly conserved network of neuron and glial cells located throughout the intestinal tract [143]. In this study, it likely was indicated in the development of gut inflammation and dysbiosis in response to psychological stressors [144], such as pain-induced anxiety [145], and could also underlie the presence of the allostatic tissue response [146]. Indeed, our preliminary data suggest a likely neurobiological involvement in the intestinal response to joint inflammation induced by CFA-IAI (*data not shown*). Additional mining of our transcriptomic data sets indicated modulation of serotonergic pathways, glutamatergic synapse activity, synaptic vesicle recycling, and retrograde endocannabinoid signaling throughout both ileum and colon. Select transcripts of genes with known neurobiological effects, such *Gdf15*, *Atp1a3*, and *Mdga2* [147] were activated in the ileum [147–149], suggesting that in future mechanistic studies for joint-gut communications, potential Vagus/ENS responses should be assessed.

In conclusion, since the CFA-induced joint inflammation model does not develop chronic systemic inflammatory conditions, it remains to be established whether multiple or chronic or consecutive allostatic responses of the intestinal tissue to joint inflammation will result in ‘allostatic overload’ [150]. Such conditions might ultimately lead to pathological disruption of the intestinal barrier function with dysbiosis as reported in both inflammatory bowel disease [151] and might also underlie the intestinal dysfunction reported in both patients and animal models of inflammatory arthritis [2, 21].

## Methods

### Animal husbandry

All procedures used in this study were approved and in compliance with the Institutional Animal Care and Use Committee (IACUC) at Rush University Medical Center. A total of 44 male Sprague Dawley (SD) rats (375-400g, approximately 4 months old, Envigo) were pair-housed in cages (polysulfone,19 × 10.5 × 8 in.) in the animal facility maintained at 22°C and 35%-55% humidity with a 12-hour light/dark cycle (light 7 AM-7 PM). Standard laboratory-grade rodent chow (Teklad Global Rodent Diet 2018, 18% protein; Envigo) and water were provided *ad libitum* (**Fig. S1**).

### Induction of knee joint inflammation with Complete Freund’s adjuvant

Experiments were initiated after a 2-week acclimatization period to minimize effects of stress conditions during transport and housing, severity, and inter-group inconsistency in disease responses. Intra-articular injections (IAI), blood collections and sacrifice were all conducted at the same time of day, i.e. between 9AM-noon. Rats were assigned to one of three study groups (n=6 rats/group): Naïve (no intra-articular injection), Complete Freund’s Adjuvant (CFA, 10mg Mycobacterium tuberculosis, 1.5ml Mannide monooleate and 8.5ml paraffin oil, InvivoGen) and vehicle controls (1xPBS). IAI was administered using a 29G insulin syringe lateral to the patellar tendon as a single 50 µl dose bilaterally, into the left and right knee joints. On day 0 (day before IAI) and days 3, 7 and 14 post-IAI, under 3% isoflurane anesthesia, rats were weighed and knee and ankle diameters measured using calipers in the medial-lateral orientation. Non-fasted tail-vein blood was collected into a BD Vacutainer SST tube per rat. Blood was allowed to clot at room temperature. Serum was separated by centrifugation at 3,000 rpm for 18 min at 4°C (∼250 µL per bleed) and 40 µL 10X proteinase inhibitor (PI) cocktail (cOmplete, Mini, EDTA-free Protease Inhibitor Cocktail) was added prior to storage at -80°C.

### Sacrifice, tissue collections and sample storage

All rats were euthanized by CO_2_ inhalation and secondary cervical dislocation. At sacrifice, bilateral hind limbs were collected from all rats and disarticulated at the hip. From right hind limbs, synovial fluid was collected by opening the joint capsule while the knee was carefully flexed. The synovial fluid from the right and left knee joints was combined from each individual rat. The joint cavity was rinsed twice with 150µL 1xPBS and collected into sterile microcentrifuge tubes. Lavages were centrifuged at 3,000 rpm/805 g for 10 min at 4°C to remove cells and debris. Supernatants were stored at -80°C until further analyses. To minimize difficulties in result interpretation due to potential sampling errors of synovium from the rat knee joint, synovial tissues were collected from the readily identifiable peri-meniscal sites and the patellar fat pad, placed into RNALater and stored at -80°C for RNA-Seq analyses. From the right disarticulated hind limbs, femora and tibiae were fixed in 10%NBF for 7 days and then moved to 70% EtOH until they underwent micro-computed tomography scanning (Scanco 50, Switzerland). Left intact knee joints were isolated by making transverse cuts 1.5cm proximal and distal to the joint and then fixed in 10% neutral-buffered formalin (NBF). Joints were decalcified in 20% EDTA, cut at midline sagittally and paraffin processed, embedded and sectioned at 6µm slices for histology. Right intact knee joints were dissected open using a fresh scalpel. Synovium and patellar fat pad were excised into RNALater (Invitrogen) and froze at -80°C until RNA extraction. Ileum and colon were dissected and fecal contents expressed into sterile 1.5ml microcentrifuge tubes from both gut segments and stored at -80°C for 16S microbiome analyses. The remaining tissues were either rinsed in ice-cold 1xPBS, transferred to RNALater and stored at -80°C for RNASeq, or placed into Modified Bouin’s fixative and prepared into Swiss rolls [152] for paraffin embedding, and sectioned at 6µm thickness for histology and immunohistochemistry (**Fig. S1**).

### Histological and immunohistochemical evaluation of joint and intestinal tissues

Sagittal sections from knee joints were stained with Hematoxylin/Eosin, anti-CD68 (Abcam, Ab283654; 1:100) or anti-CD4 Abcam, Ab237722 1:2000). Sections of intestinal Swiss rolls were stained using the Alcian Blue Kit (Vector Labs, H-3501), anti-CD68 (1:100), anti-CD4 (1:2000), anti-CD8 (Abcam, Ab237709, 1:500) or anti-Ki67 (Invitrogen, MA5-14520; 1:100), Invitrogen, MA5-14520) or the following biotinylated lectins (all from Vector Labs), *Griffonia Bandeiraea Simplidifolia* Lectin I, (GSL I, B-1105-2), *Maackia Amurensis Lectin II* (MAL II, B-1265-1) and *Ulex Europaeus Agglutinin I* (UEA, B-1065-2). Prior to histochemical staining, antigen retrieval was performed using Tris/EDTA (Abcam, ab93684), and sections were blocked with normal goat serum (Vector Laboratories, S-1000). Immuno- and lectin reactivities were developed using the ABC kit (Vector Laboratories, PK-7100) and the DAB kit (Vector Laboratories, SK-4100). Methyl green (Vector Laboratories, H-3402) was used as a counterstain. Slides were imaged using an Olympus VS200 Scanner.

### Western blot characterization of affinity-purified antibodies to mouse bikunin

A synthetic peptide of mouse bikunin was synthesized and conjugated to Keyhole Limpet Hemocyanin (KLH), via a non-authentic cysteine residue at the C-terminal end, for production of a polyclonal antibody in a rabbit (Mimotopes, Australia). The anti-serum (from rabbit A) was affinity purified using the immunizing peptide (Genscript, USA) covalently coupled via the cysteine residue to SulphoLink Coupling Resin (Thermo Scientific Pierce, UK), according to the manufacturer’s instructions. Purified IgG fractions were pooled, concentrated and stored in aliquots at -20°C; the affinity purified antibody is designated ap_A_mBikunin. A Western blot of mouse serum treated with/without chondrotinase ABC lyase or NaOH was used to characterize the affinity purified antibodies. Here 1.25 µL of mouse serum was incubated with/without 0.1 U Chondroitinase ABC lyase (from *Proteus vulgaris*; Sigma Aldrich) for 2 h at 37°C or 0.1 M NaOH for 10 min at RT after which 1.0 μL 1 M HCl was added to the NaOH-treated samples. Reactions were stopped by adding 5 µl of 5X sample loading buffer; 12 µl of each reaction was run on a NOVEX 4-20% Tris-Glycine gel. Proteins were transferred to a nitrocellulose membrane in Tris-Glycine transfer buffer. Blots were blocked for 1 h with 5% (w/v) dried milk in PBS/0.05% (v/v) Tween-20 (PBST), washed briefly in PBST and then incubated with ap_A_mBikunin (diluted 1:500 in PBST with 5% (w/v) dried milk) overnight at 4°C. After washing in PBST the membrane was incubated with IRDye 800CW-conjugated goat anti-rabbit IgG secondary antibody (1:5000 in PBS; LI-COR Biosciences Ltd) for 1 h at room temperature, washed as before and visualized using an Odyssey CLx imaging system (LI-COR). The species detected by the antibody are indicated to the right in Fig. S2.

### Western blot analyses of serum and synovial fluid bikunin

Commercially available mouse serum, which was processed the same as experimental rat serum, was included in each western blot as an internal standard. Serum samples were thawed on ice and diluted 1:1 with 50mM ammonium acetate pH 7.0 containing 1XPI and debris removed with a 0.45µM filters. 150µL of sera were desalted using MicroCon 3 filtration units (at 12,500 rpm in a microfuge for 15 min) and retentates washed once with 450µL 50mM ammonium acetate, pH 7.0. Desalted proteins were collected from the membrane with 500µL ammonium acetate, pH 7.0. Portions (containing 7.5µL equivalents of serum) were incubated at 37°C for 2 hrs. with or without 5mU of protease-free chondroitinase ABC lyase (Chondroitinase ABC protease free).

Synovial fluid samples were thawed on ice, diluted to 450µL with 50mM ammonium acetate, pH 7.0 containing 1XPI, clarified by centrifugation at 12,000 g at 4°C for 15 min, and desalted using MicroCon 3 filtration units as described above for serum samples. The retentates were recovered in 200µL 50mM ammonium acetate, pH 7.0 and 100µL portions were incubated at 37°C for 2h with or without 5mU of protease-free chondroitinase ABC.

Sera and synovial fluid samples were speedvac dried and analyzed by gel electrophoresis and western blot. Chondroitinase digestion buffer was removed by speedvac evaporation. Samples were dissolved in Tris-glycine SDS sample buffer (Novex, AMS.E1028-02) containing 100µl of 0.5mM DTT, heated at 90°C for 5 min before electrophoresis on SDS gels (Invitrogen Novex WedgeWell 4-12% Tris-Glycine). Separated proteins were electro-transferred on ice (250V, 302µA for 45 min and 150V, 297µA for 15 min) to 0.45µm nitrocellulose membranes (BioRad) and these were incubated with affinity purified anti-bikunin antibody (ap_A_mBikunin). Antibody reactivity with inter-α-inhibitor (IαI) and pre-α-inhibitor (PαI), bikunin, and bikunin core protein[108] was characterized (**Fig. S2A).** The Western blotting procedure (in **Fig. S2 B,C**) was performed as previously published [153]. Chemiluminescent images were recorded using iBright1500 Instrumentation and further analyzed using ImageJ software. Data are shown as integrated pixel density (IPD) relative to bikunin present in standard mouse serum (Sigma Aldrich m595).

### Micro-computed tomography analyses of femurs

Micro CT scanning was completed using a Scanco μCT50 (Switzerland). Right femurs were scanned using 70kVp, 114μA, 500ms integration time, and 10μm voxel size. The region of interest (ROI) for trabecular and cortical bone analyses started just proximal to the distal femoral growth plate and continued 2 mm proximally into the metaphysis. The trabecular and cortical bone compartments were analyzed separately using thresholds of 200 and 1000, respectively. Cortical and trabecular bone parameters were measured using the manufacturer’s software. Periosteal reaction was noted when present and analyzed separately from the cortex using the same threshold values.

### Bulk RNASeq analyses of synovium and intestine

To obtain reproducible and high-quality yields of RNA from synovium, tissues were harvested from right and left knees and pooled from n=3 rats, then analyzed as a single sample. Distal ileum and colon tissues in RNAlater were thawed on ice and 1cm pieces were dissected out (including villi, submucosa, and smooth muscle layers) using transverse cuts. The number of intestinal tissue replicas to provide sufficient DEG detection power was determined as previously described [154].

Using previously published methods [155] RNA was extracted and purified from all tissues, samples with RIN numbers ≥7 were reverse transcribed into cDNA, PCR amplified with primers CS1_515F and CS2_806R followed by library preparation. Bioinformatic analyses of sequencing data was carried out as follows: Raw reads were trimmed to remove Truseq adapters and bases from the 3’ end with quality scores less than 20 using cutadapt[156]; trimmed reads shorter than 40bp were discarded, and were then aligned to the mRatBN7.2 *Rattus norvegicus* (Norway rat) reference genome using STAR[157]. The expression level of ENSEMBL genes was quantified using featureCounts[158] and the resulting expression counts were used to compute differential expression statistics with edgeR[159, 160]. Normalized expression was computed as log_2_ counts per million (CPM), including a TMM normalization and batch effect correction. In all cases p-values were adjusted for multiple testing using the false discovery rate (FDR) correction of Benjamini and Hochberg[161]. Volcano plots were generated within the R programming language. DEG (Differentially Expressed Genes) with an absolute Log_2_FC (Log Fold Change) <+2 or <-2 and a q value of less than 0.05 were assigned to specific pathways using the KEGG and further grouped under seven functional categories: Metabolism, Cellular Processes, Genetic Information Processing, Environmental Processing and Immune System

### 16S microbiome analyses

Details have been described by us previously [6]. Briefly, DNA was extracted from fecal samples using a Maxwell RSC48 device (Promega), with bead-beating implemented off-instrument. Microbial community structure was characterized by DNA extraction, PCR amplification and sequencing of 16S ribosomal RNA gene amplicons[162]. Microbiome bioinformatics was performed with QIIME2 2021.11[163]. Raw sequence data were checked for quality using FastQC and merged using PEAR[164]. Merged sequences were quality filtered using the q2-demux plugin followed by denoising with DADA2 [165] (via q2-dada2). Primer adapter sequences were removed using cutadapt algorithm[156]. Alpha-diversity metrics (observed features[166], Shannon Index[167], Simpson’s Index [168], and Pielou’s Evenness) and beta diversity metrics were calculated using q2-diversity after samples were rarefied to a depth of 59,000 sequences per sample. Taxonomy was assigned to ASVs using the q2-feature-classifier[169] classify-sklearn naïve Bayes taxonomy classifier against the SILVA 138 99% reference sequences database[170]. Differentially abundant genus level taxa between pairwise groups were identified using the compositional centered Log-ratio Kruskal Wallis (CLR-KW) algorithm. Adjusted P-values (also known as q-values) were generated using the Benjamini–Hochberg method[161].

### Statistical Analyses for Bone Data

Control and test rats from 2 separate cohorts (one in the summer and one in the winter) were included. For the trabecular parameters, we observed a cohort effect so we express all test results normalized to the mean of the cohort-specific control animals. Significance was assessed using one-way analyses of variance (ANOVA) for group effects with Bonferroni post hoc tests. All results are reported as means ± standard deviation and individual data points are shown. The statistical testing was performed with commercially available software (SPSS v.19 for Windows, Chicago, IL).

## Acknowledgements

We would like to thank Dr. Georgia Papavasiliou (Illinois Institute of Technology) and Dr. Joseph Reynold (Rosalind Franklin University) for their insightful discussions on intestinal immunology and therapeutic target discovery during the progression of this project; Dr. Stefan Green for his input in study design and genomic analyses; and Rylan Martin for technical support. Funding for this project is from the Orthopedics Departmental Fund RUMC (MMM), the Katz/Rubschlager Endowed Chair at RUMC (AP), and Versus Arthritis, UK Grant 22277 (AJD and CMM).

## Supporting Information Figures

**Supporting Information Fig. S1.**
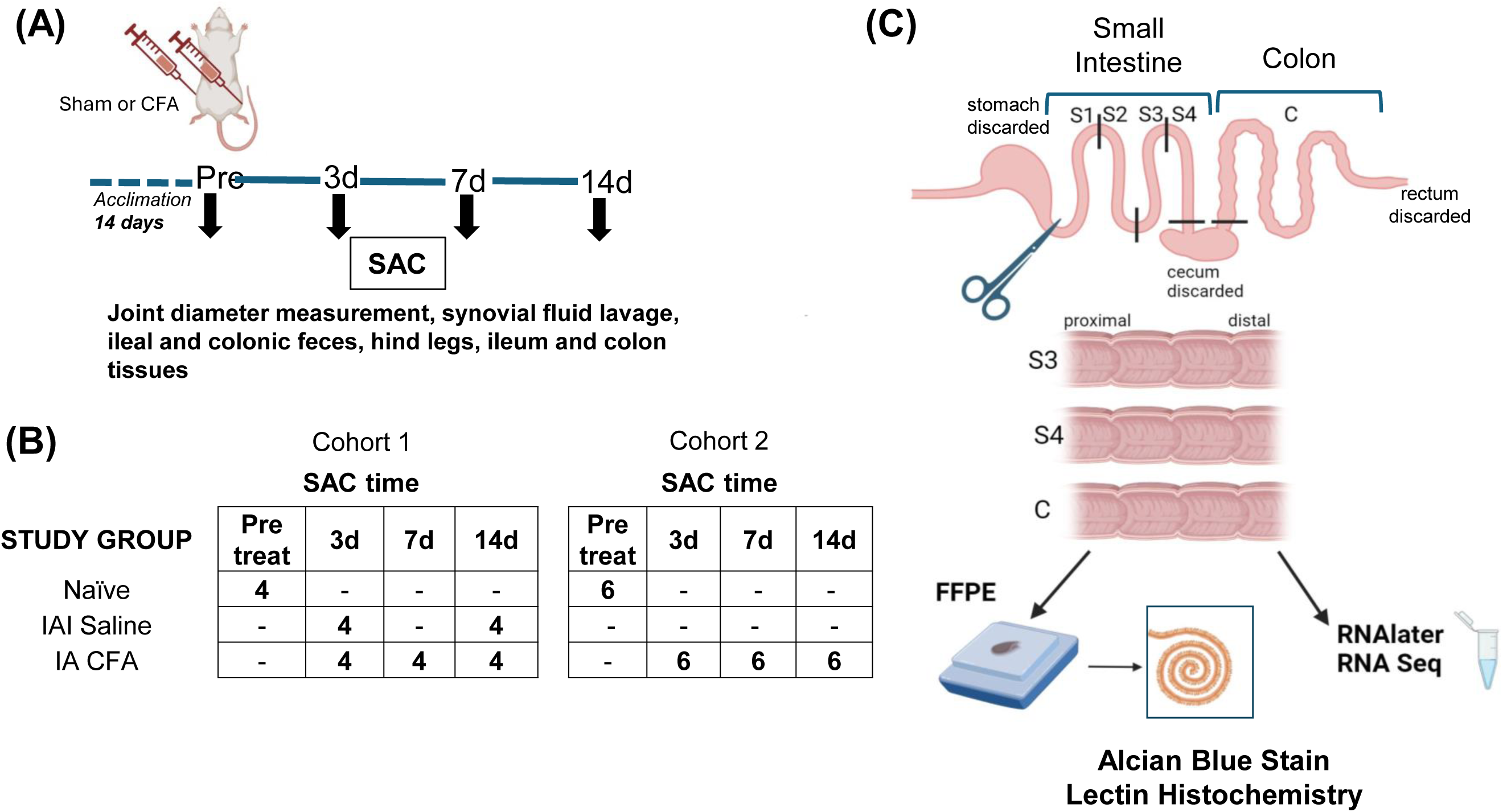

**Supplemental Figure S2.**
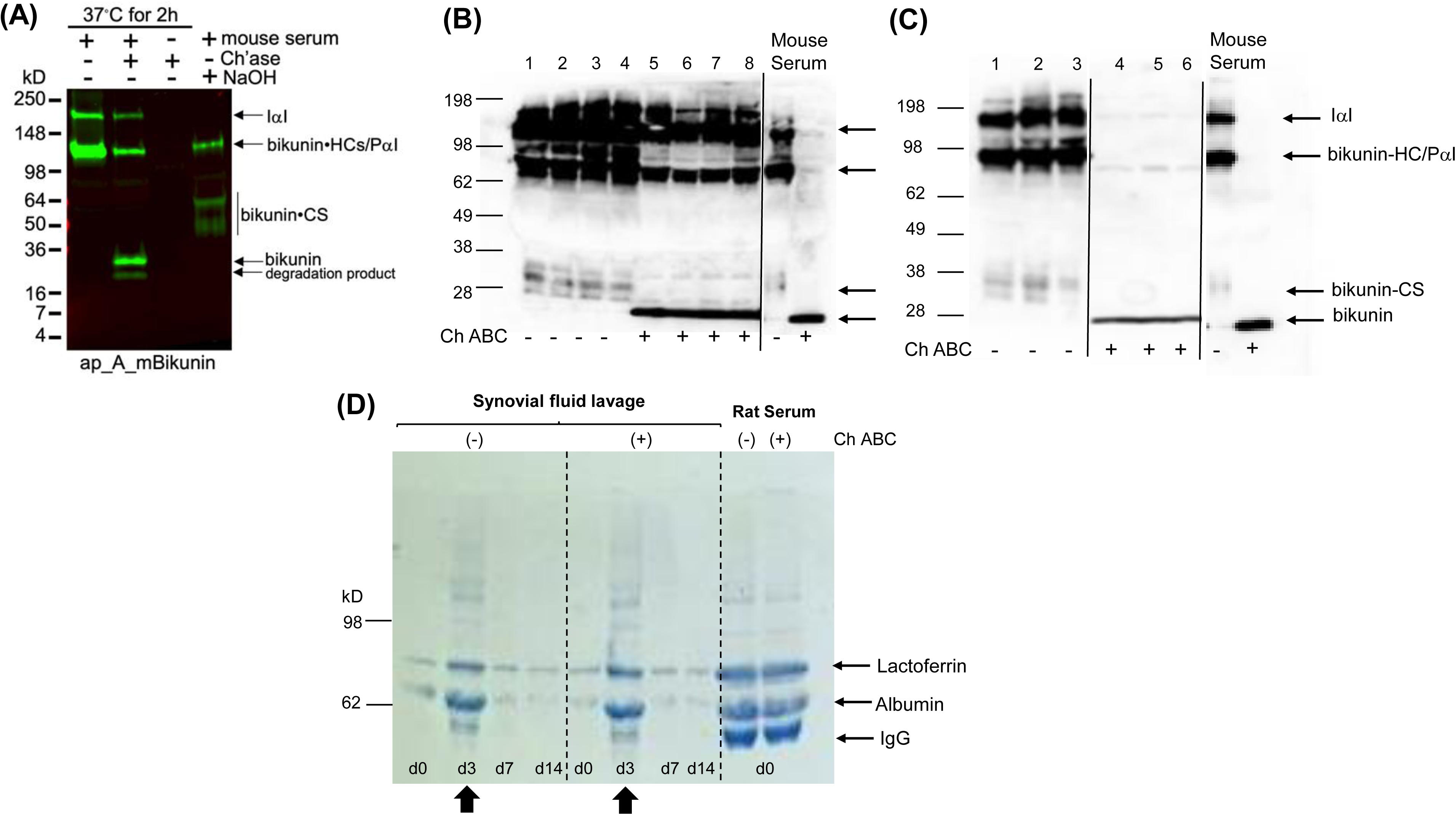

**Supporting Information Fig. S3.**
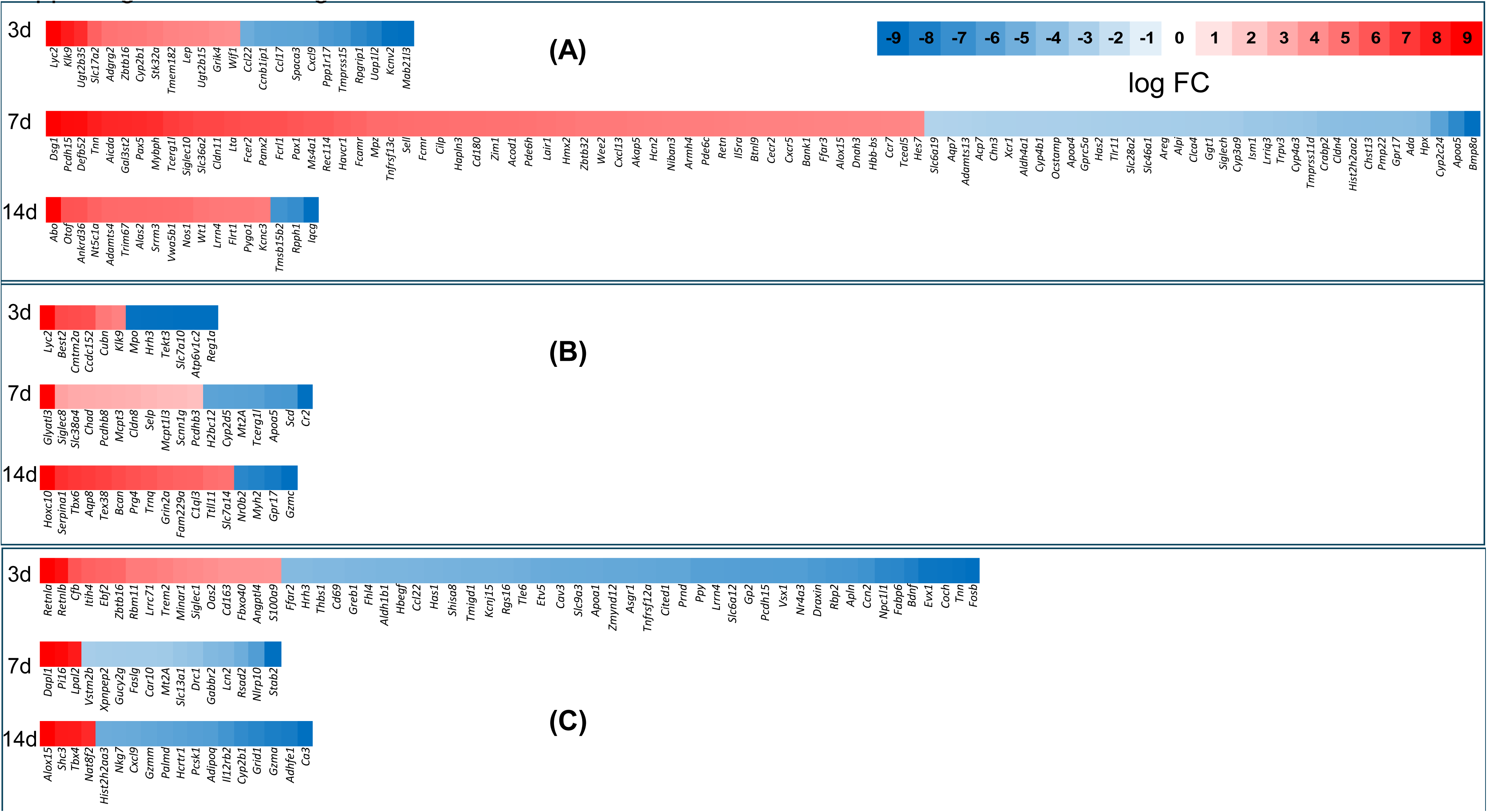

**Supporting Information Fig. S4.**
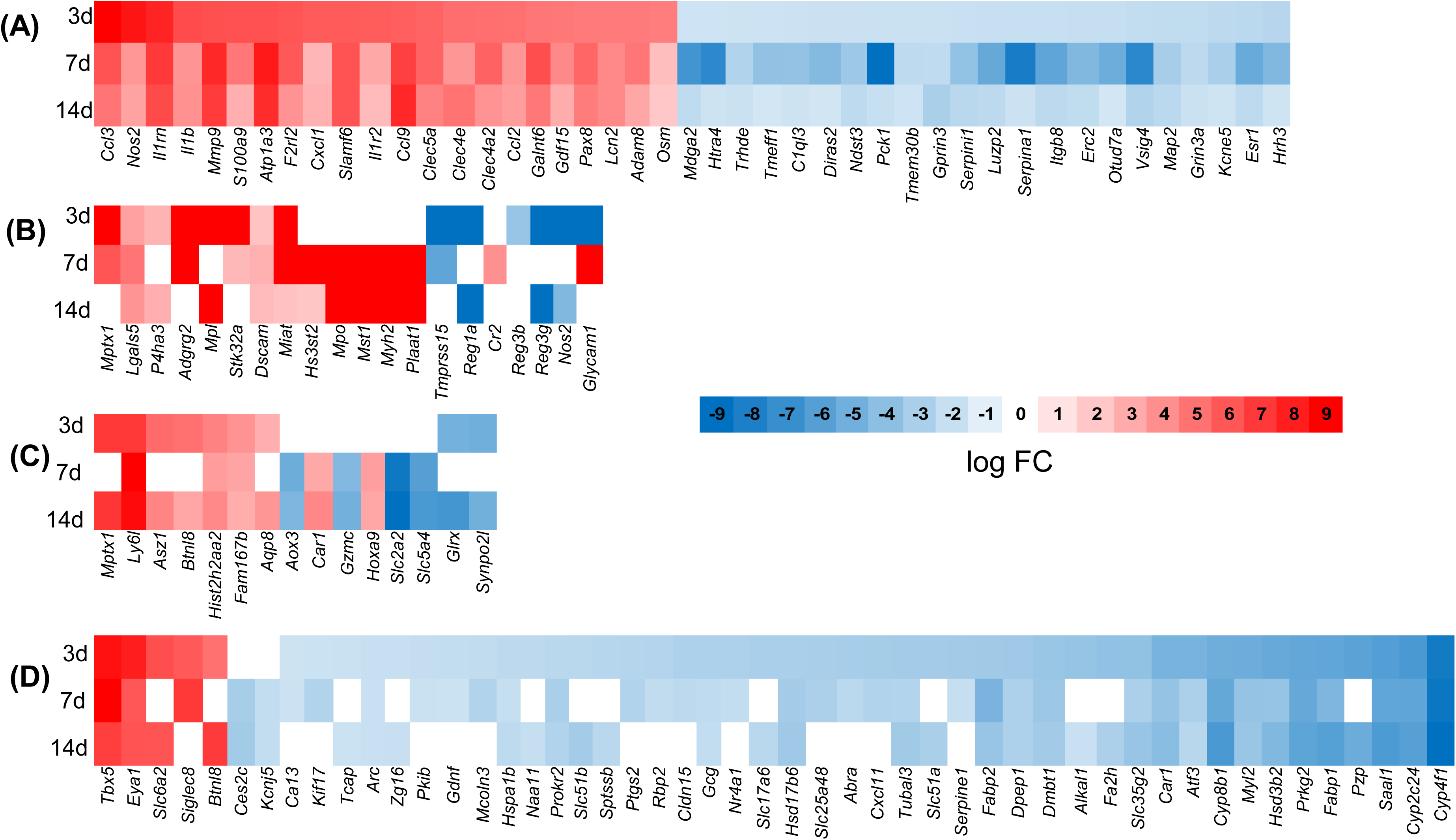

**Supporting Information Fig. S5.**
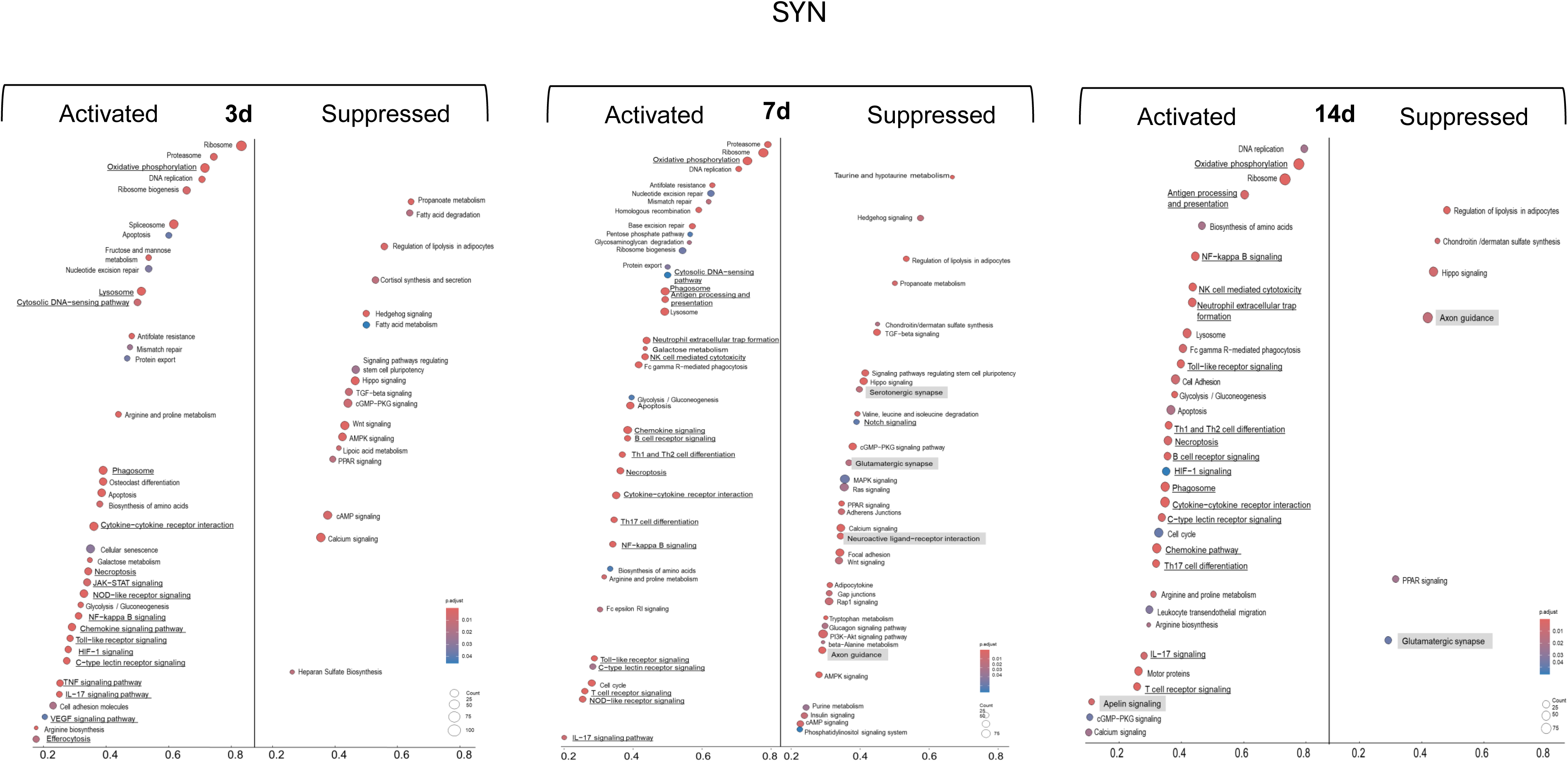

**Supporting Information Fig. S6.**
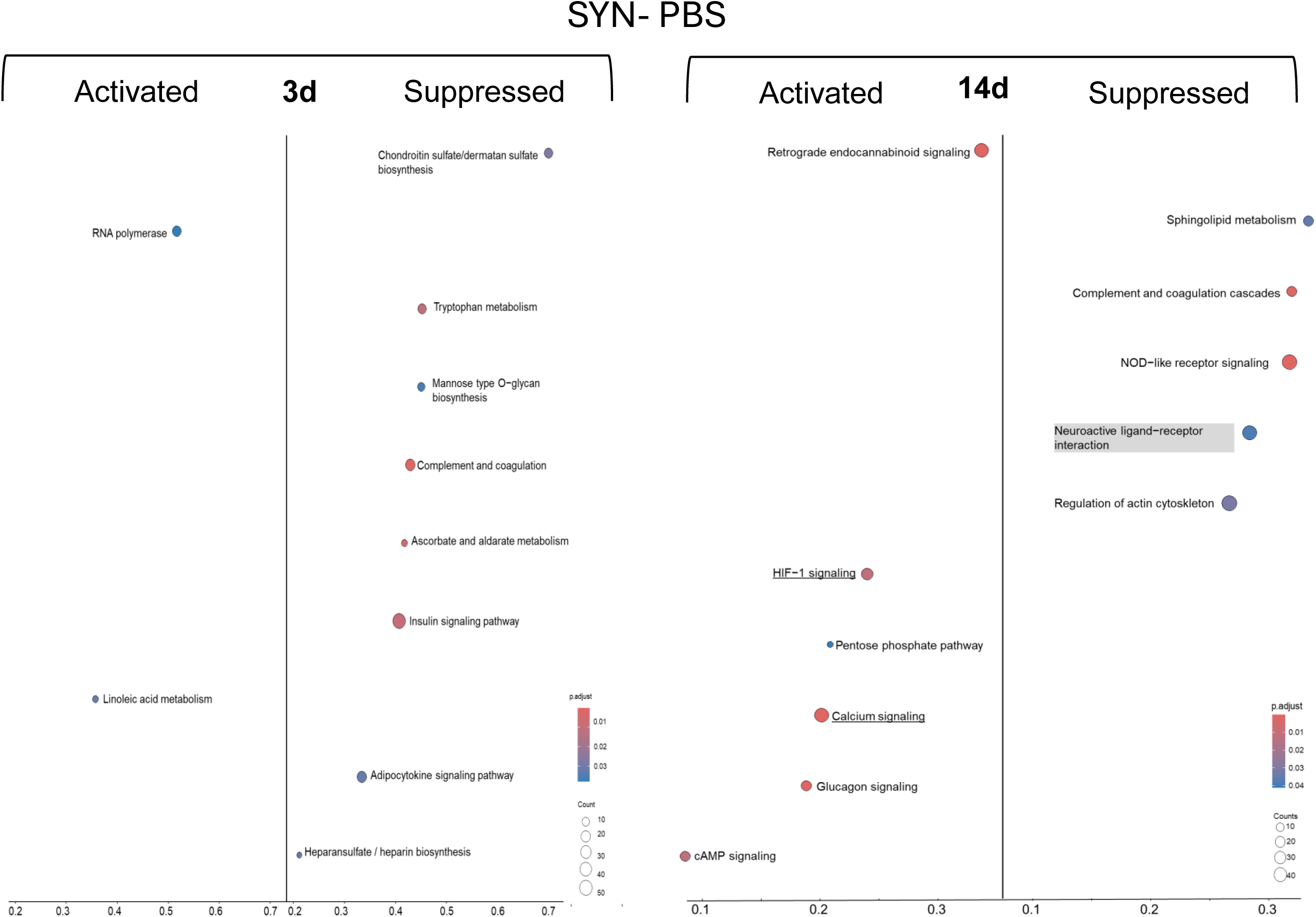

**Supporting Information Fig. S7.**
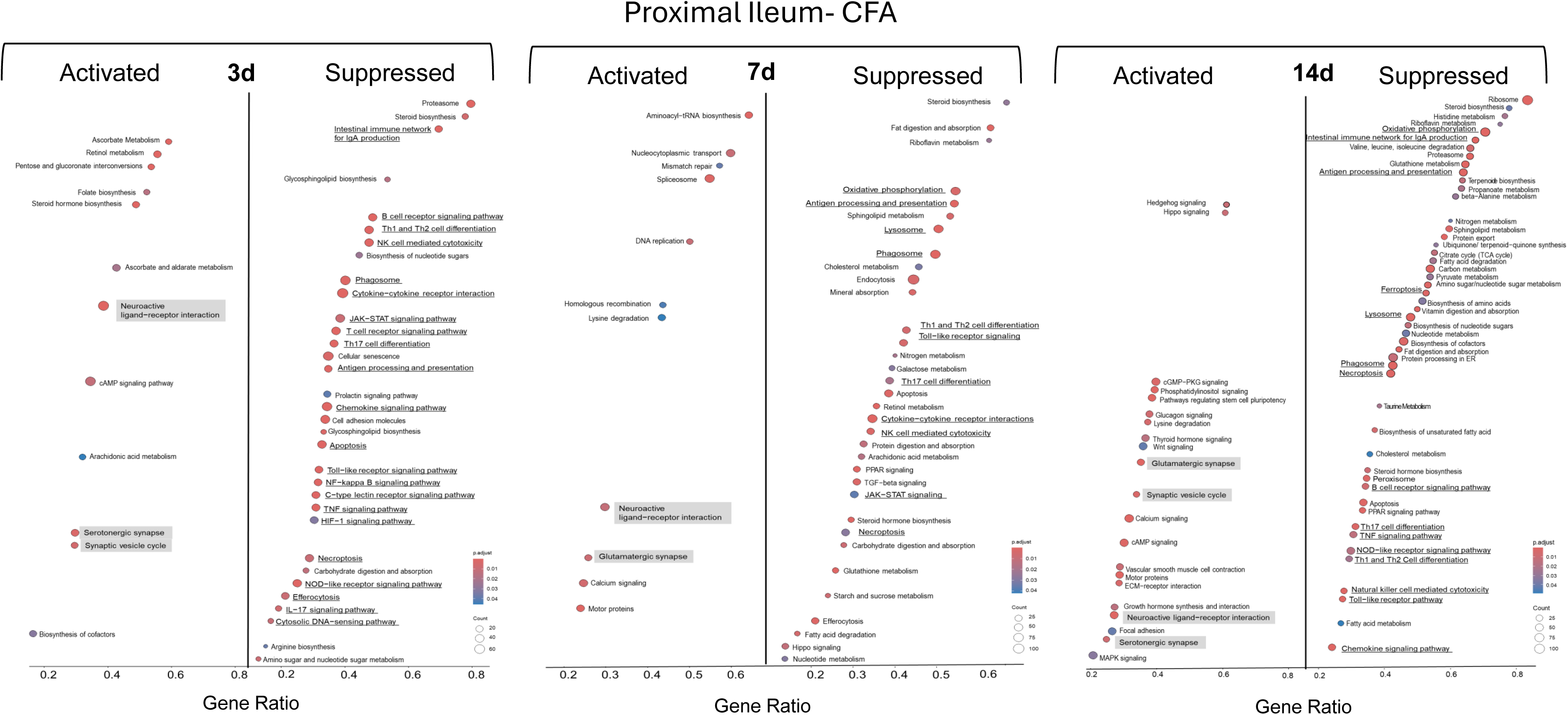

**Supporting Information Fig. S8.**
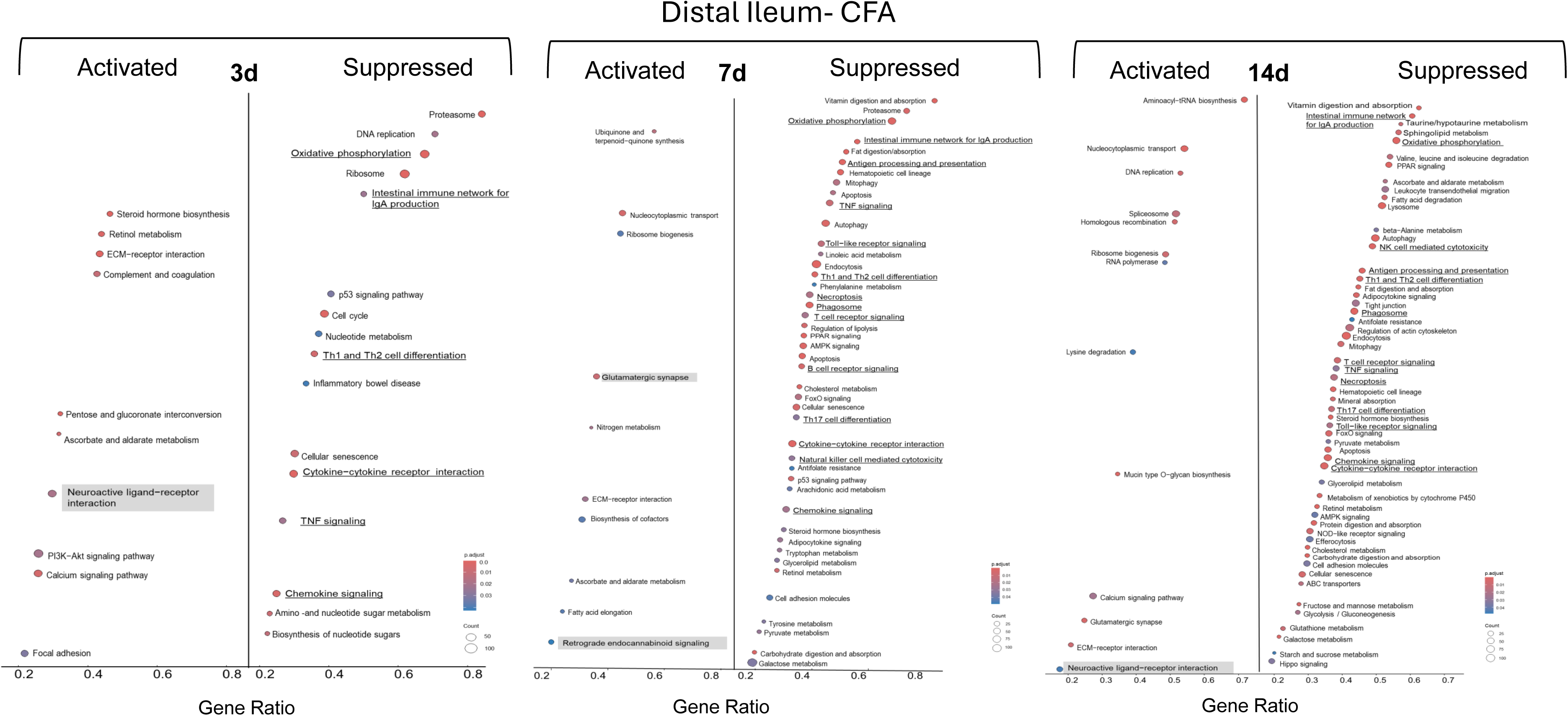

**Supporting Information Fig. S9.**
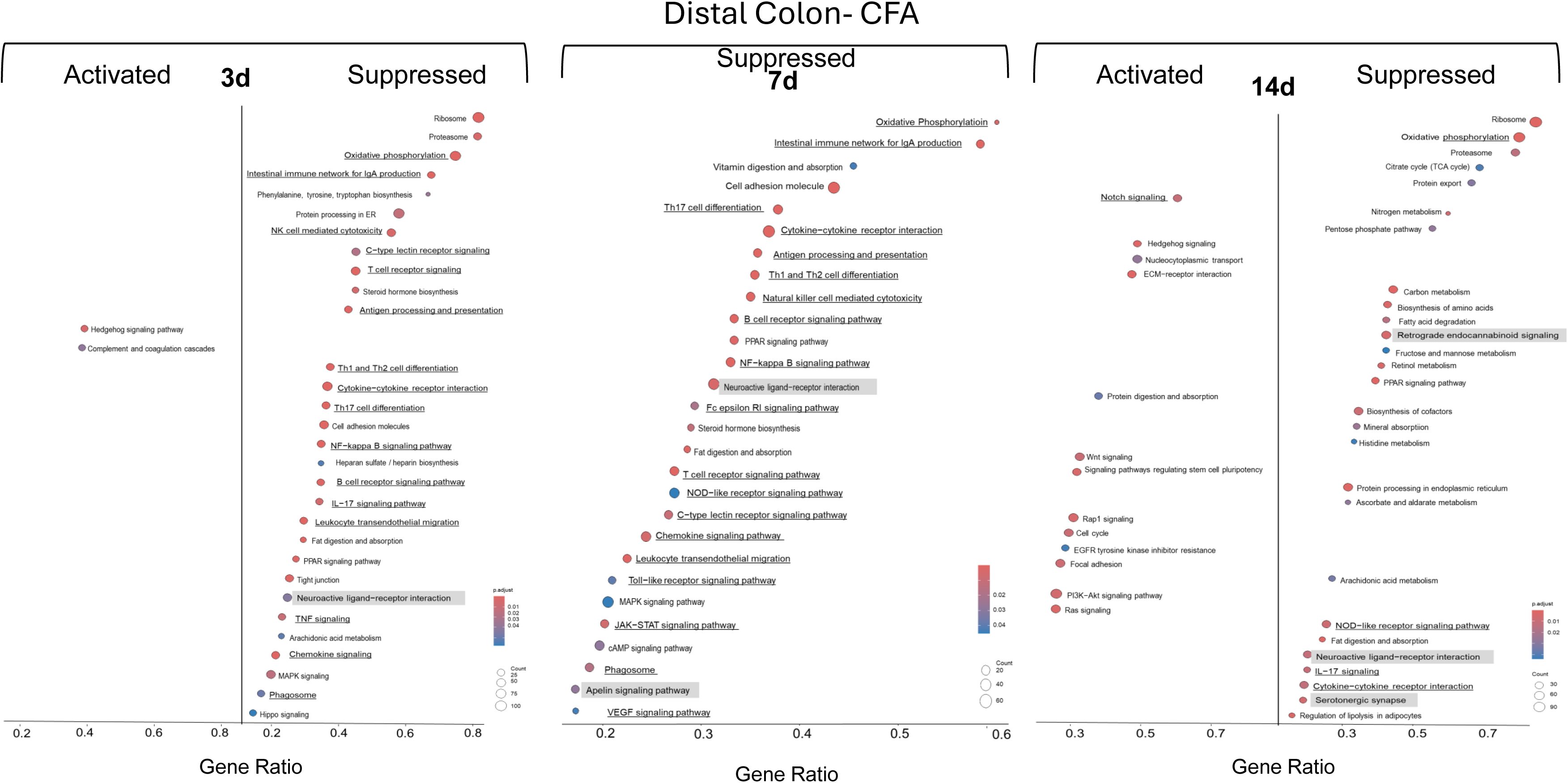

**Supporting Information Fig. S10.**
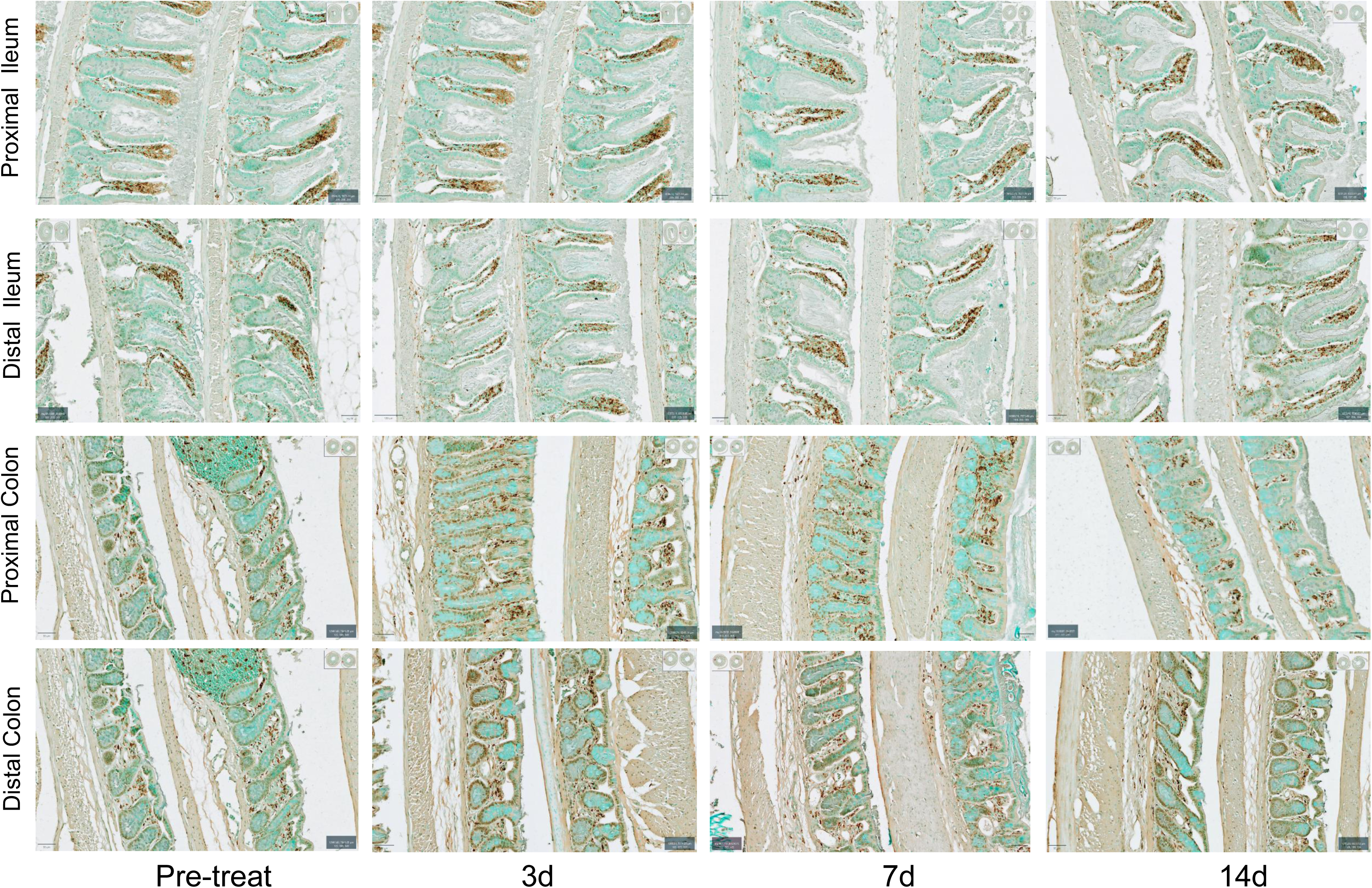

**Supporting Information Fig. S11.**
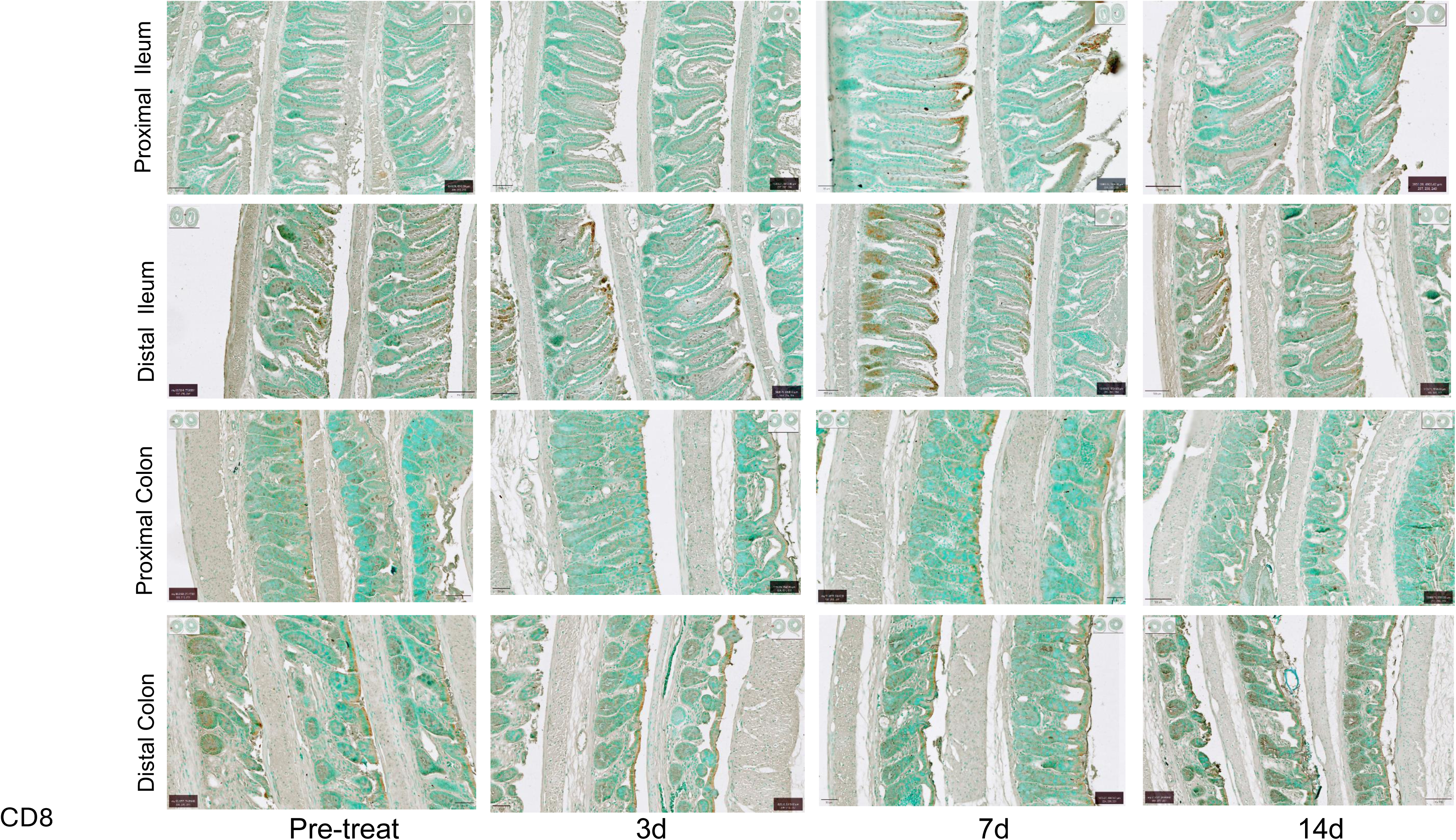

**Supporting Information Fig. S12.**
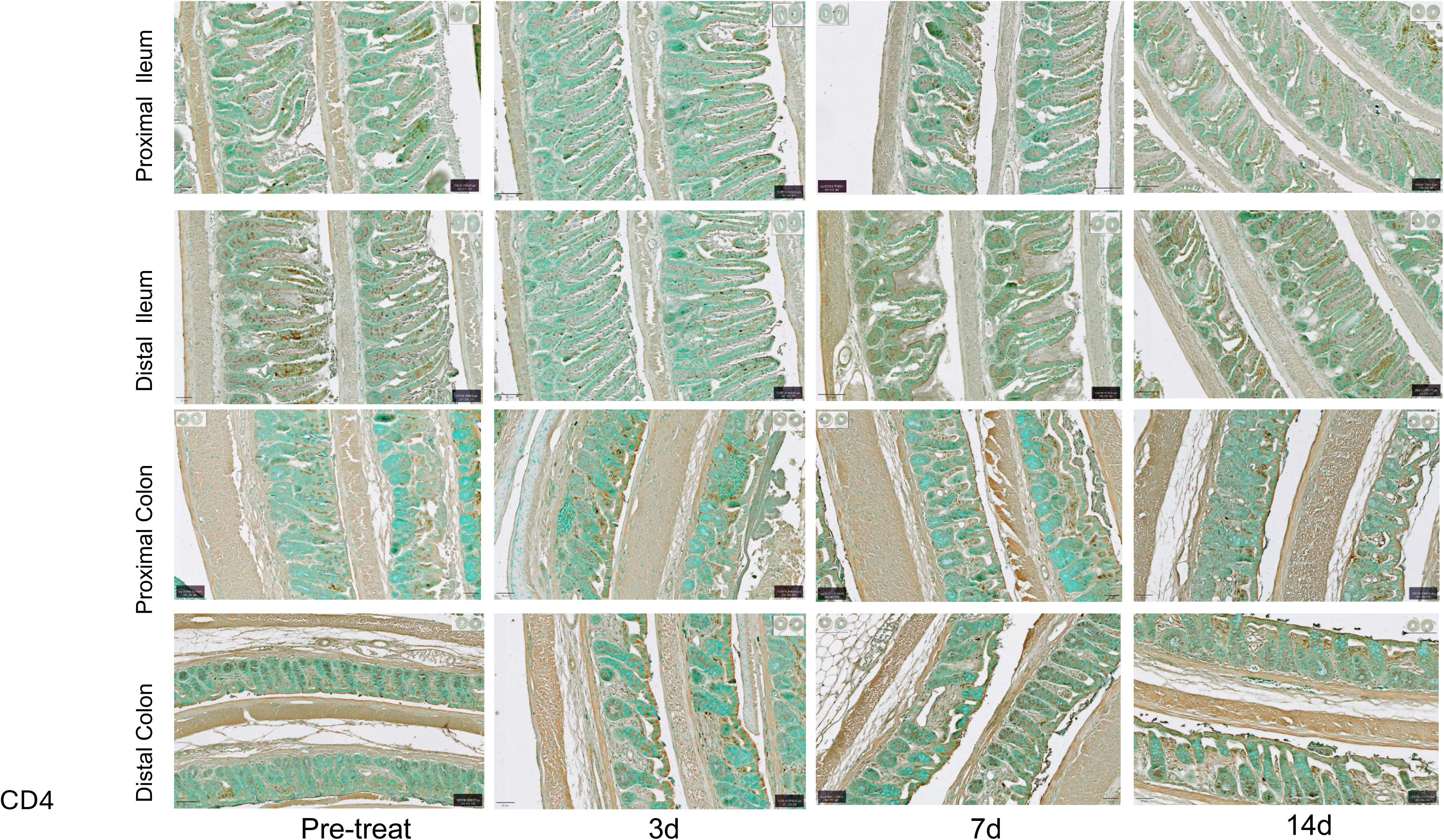

**Supporting Information Fig. S13.**
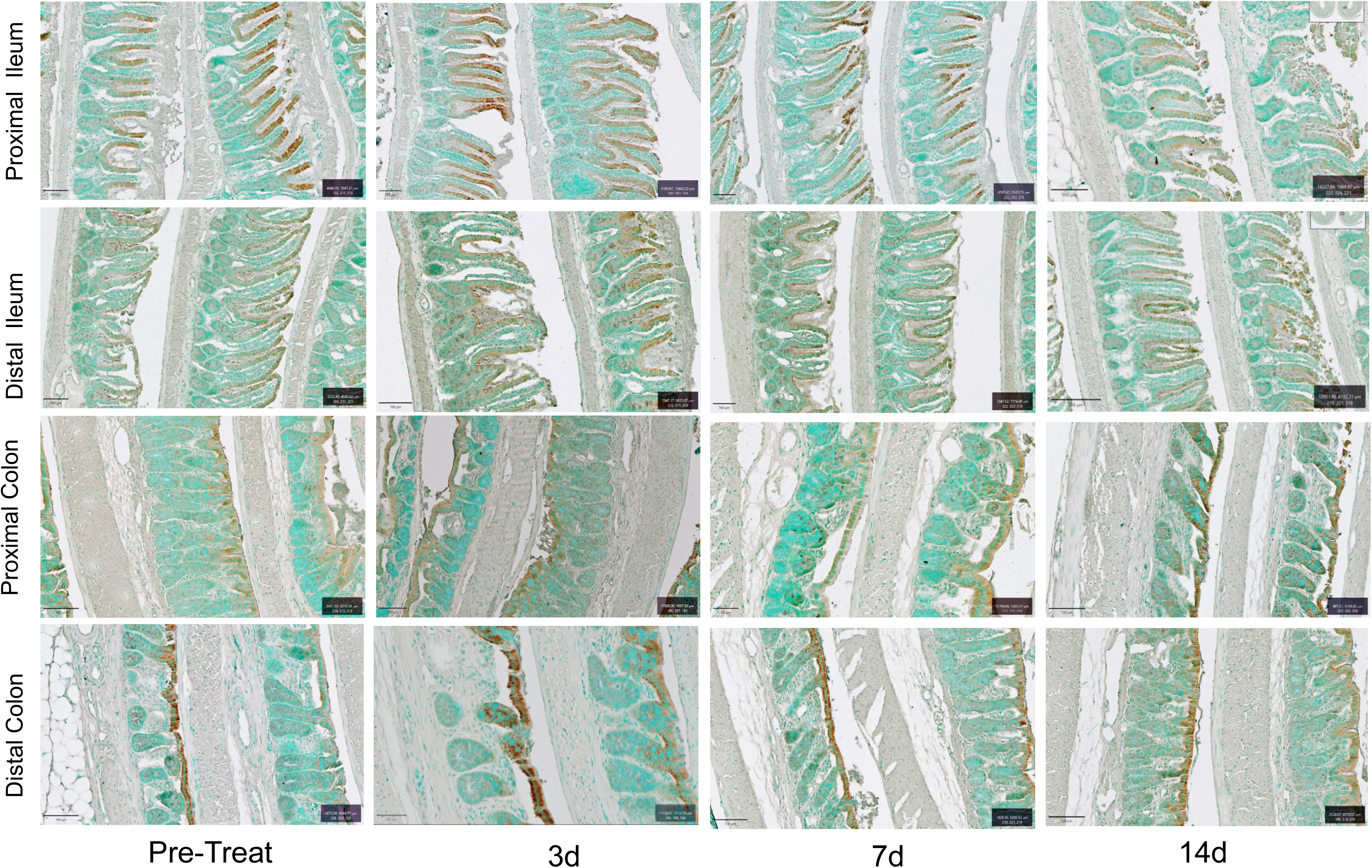

**Supporting Information Fig. S14.**
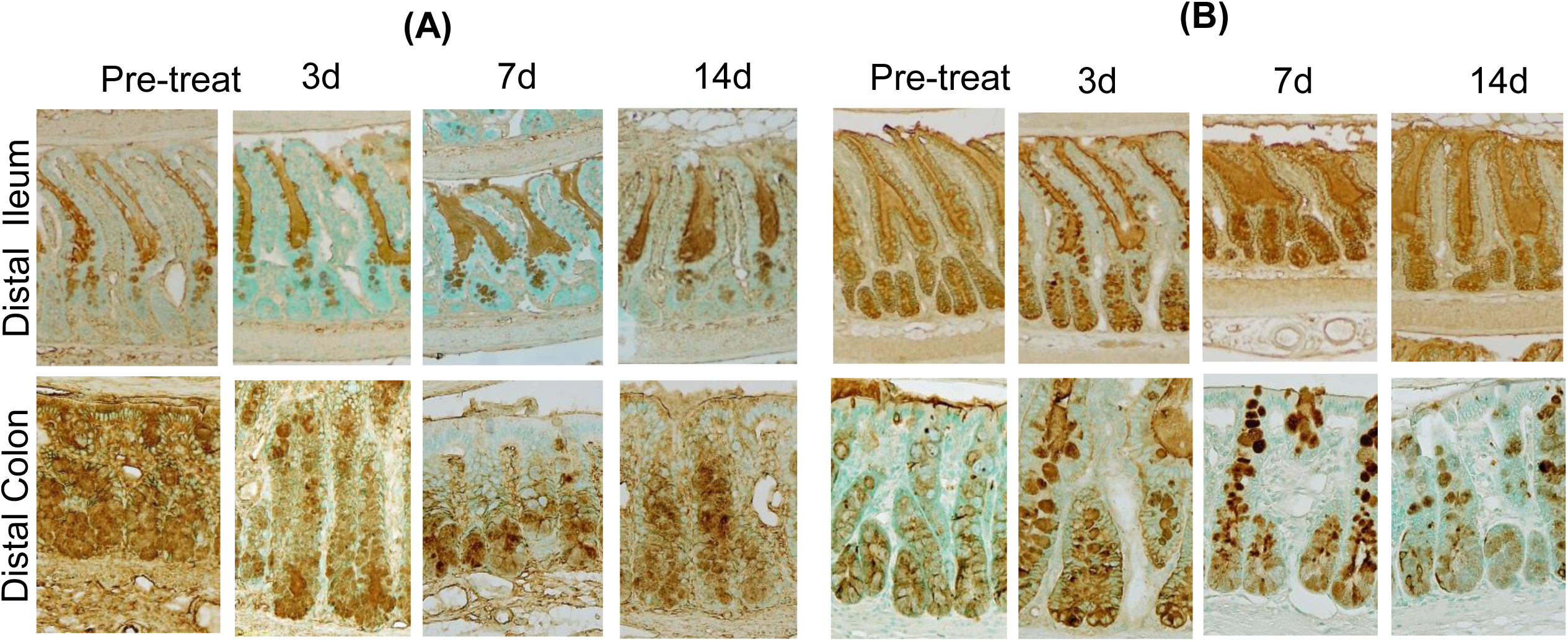

**Supporting Information Fig. S15.**
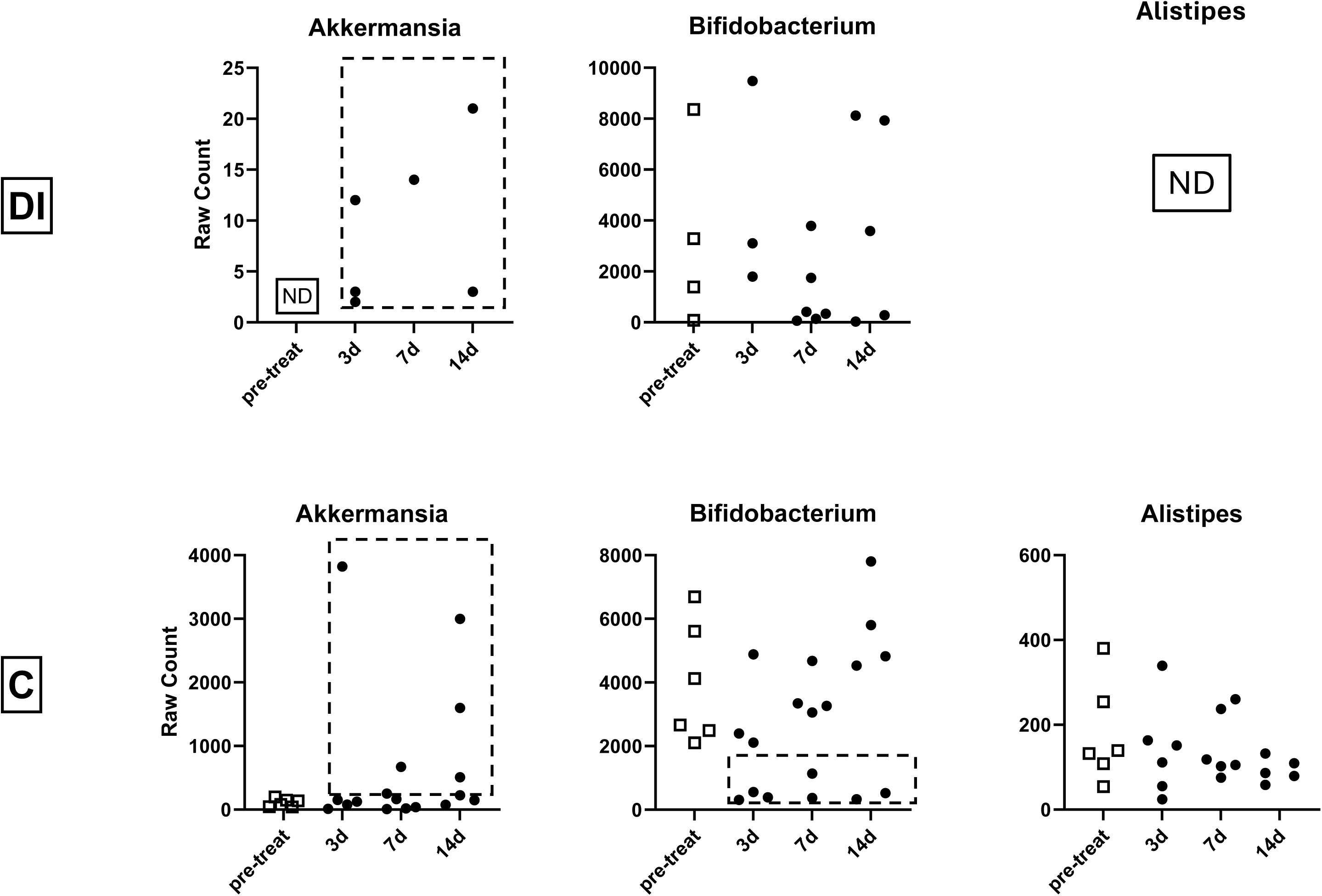

