## Supplemental Figure legends for "Evidence of an Allostatic Response by Intestinal Tissues Following Induction of Joint Inflammation"

***Supplemental Figure S1.*** **Overview of experimental protocol.**

**(A)** Schematic of *in vivo* CFA Model (see Methods for Details). Rats received one intra-articular injection (IAI) of 50 µL of CFA (10mg/ml) or sterile saline into both knee joints **(B).** Number of rats used for each study group (Naïve, Saline, CFA). Experiments were performed on 2 separate cohorts, but procured from the same supplier. S3= Proximal Ileum (PI), S4=Distal Ileum (DI) C=Colon (Co). **(C)** Segmentation of small intestine and colon for RNASeq analyses and histological evaluations (see Methods for details). S=small intestine, S3=proximal Ileum, S4= distal Ileum, C= distal colon

***Supplemental Figure S2***. **Characterization of anti-bikunin antibody ap_A_mBikunin, Western blots analyses of bikunin complexes in rat sera and synovial fluid, and detection of serum proteins in knee joint synovial fluids post-IAI-CFA.**

**(A)** A Western Blot of mouse serum treated with/without chondroitinase ABC lyase (Ch'ase) or NaOH to release bikunin or bikunin•CS, respectively from IαI, PαI and bikunin•HS species was used to characterize a rabbit anti-mouse bikunin antibody (denoted ap_A_mBikunin) as described in the Methods. Note that the mouse sequence of the immunizing peptide, AVLPQE**S**EGS, is highly similar to the corresponding rat sequence, AVLPQE**N**EGS, with only one amino acid difference. **(B)** Western blot of rat sera, collected from 4 individual rats prior to IAI-CFA, and mouse serum (as a standard) were prepared and treated without (lanes 1-4) or with (lanes 5-8) chondroitinase ABC lyase (Ch ABC) prior to electrophoresis as described in the Methods. **(C)** Western blot of rat sera of knee joint synovial fluid lavages collected from 3 individual rats at d3 post IAI-CFA and 'standard' mouse serum were prepared and treated or not with chondroitinase ABC (Ch ABC) prior to electrophoresis as described in the Methods. Positions of bikunin-contianing species in **B** and **C** (i.e., IαI, PαI, bikunin-CS and bikunin) are indicated to the right of the blots. All membranes (in **A**, **B**, **C**) were incubated with ap_A_mBikunin as described in the Methods. **(D)** To assess effusion of serum proteins into the joint space following IAI-CFA, portions of rat synovial fluid lavages collected prior to IAI-CFA (d0) and at d3, d7 and d14 post IAI-CFA, and rat serum pre-IAI-CFA, were electrophoresed with or without prior treatment with chondroitinase ABC (Ch ABC), and gels were Coomassie stained. Expected migration positions of lactoferrin, albumin and IgG are shown to the right.

***Supplemental Figure S3***. **Heat map of DEGs in proximal ileum (A), distal ileum (B) and distal colon (C) modified only at a single time point post-CFA-IAI.**

***Supplemental Figure S4*****.** **Heat map of DEGs in proximal ileum (A), distal ileum (B) and distal colon (C) modified at 2 or 3 time points post-CFA-IAI.**

***Supplemental Figure S5.*** **Dot plot representation of transcriptomic changes in synovial tissue following CFA-IAI.**

Pathways (ranked by p-value) were assigned to color-coded functional categories as listed in the KEGG Database (Metabolism, Cellular Processes, Genetic Information Processing, Environmental Processing, Immune System, Nervous System). Gene ratio for a given pathway is computed as the percentage of genes present divided by the total number of genes in that pathway. Activated and Suppressed pathways are separated by a solid black vertical line and gene ratios < or > 0.5 are separated by a dotted vertical line. The top 20 modified pathways are above the horizontal dotted line.

***Supplemental Figure S6.*** **Dot plot representation of transcriptomic changes in synovial tissue following PBS-IAI.**

Pathways (ranked by p-value) were assigned to color-coded functional categories as listed in the KEGG Database (Metabolism, Cellular Processes, Genetic Information Processing, Environmental Processing, Immune System, Nervous System). Gene ratio for a given pathway is computed as the percentage of genes present divided by the total number of genes in that pathway. Activated and Suppressed pathways are separated by a solid black vertical line and gene ratios < or > 0.5 are separated by a dotted vertical line. The top 20 modified pathways are above the horizontal dotted line.

***Supplemental Figure S7.*** **Dot plot representation of transcriptomic changes in the proximal ileum following CFA-IAI.**

Pathways (ranked by p-value) were assigned to color-coded functional categories as listed in the KEGG Database (Metabolism, Cellular Processes, Genetic Information Processing, Environmental Processing, Immune System, Nervous System). Gene ratio for a given pathway is computed as the percentage of genes present divided by the total number of genes in that pathway. Activated and Suppressed pathways are separated by a solid black vertical line and gene ratios < or > 0.5 are separated by a dotted vertical line. The top 20 modified pathways are indicated by a dotted vertical line.

***Supplemental Figure S8*.** **Dot plot representation of transcriptomic changes in the distal ileum following CFA-IAI.**
Activated pathways are listed on the left and suppressed pathways on the right. Gene ratios and pathway groupings are assigned as described for Figure S6.

***Supplemental Figure S9*.** **Dot plot representation of transcriptomic changes in the colon following CFA-IAI.**
Activated pathways are listed on the left and suppressed pathways on the right. Gene ratios and pathway groupings are assigned as described for Figure S6.

***Supplemental Figure S10*.** **Immunohistochemical localization of CD68+ cells in proximal and distal regions of the ileum and colon.**
Images represent regions from the scanned slides that were used for panels shown in Fig. 10

***Supplemental Figure S11*.** **Immunohistochemical localization of CD8+ cells in proximal and distal regions of the ileum and colon.**
Images represent regions from the scanned slides that were used for panels shown in Fig. 10

***Supplemental Figure S12*.** **Immunohistochemical localization of CD4+ cells in proximal and distal regions of the ileum and colon.**
Images represent regions from the scanned slides that were used for panels shown in Fig. 10

***Supplemental Figure S13*.** **Immunohistochemical localization of Ki67+ cells in proximal and distal regions of the ileum and colon.**
Images represent regions from the scanned slides that were used for panels shown in Fig. 10.

***Supplemental Figure S14. Lectin histochemistry in proximal and distal regions of the ileum and colon.***
Images represent regions from the scanned slides that were used for panels shown in Fig. 10. Mal II (A) and UEA (B).

***Supplemental Figure S15.*** **Abundance of mucin degrading bacteria in feces from distal ileum and distal colon pre- and post-CFA-IAI.**

Raw counts for mucin-degrading bacteria (Akkermansia, Alistipes) and sialic acid cleaving bacteria (Bifidobacterium) are shown. Note that in pre-treatment rats, Akkermansia and Alistipes were below detection in the ileum (ND= not detected). Variable animal-to-animal responses are outlined in dotted lines. Note the much lower counts for *Akkermansia* and *Alistipes* in the ileum compared to the colon. Bifidobacterium showed higher counts in both intestinal regions.
